# Anthropogenic nitrogen deposition restructures decomposing microbial communities, altering SOM molecular composition, but not molecular complexity or diversity

**DOI:** 10.1101/2025.10.21.683748

**Authors:** Brooke E. Propson, William A. Argiroff, Grace A. Cagle, Rima A. Upchurch, Donald R. Zak, A. Stuart Grandy, Zachary B. Freedman

## Abstract

Soil organic matter (SOM) consists of diverse biochemical constituents, spanning a spectrum of chemical complexity, and the relative abundance of these substrates influences microbial metabolism and soil carbon persistence. However, mechanistic controls governing these processes and how they may be affected by environmental change remains incomplete. This study aims to assess (1) the molecular-level changes that occur across stages of root decomposition, from undecayed plant root litter to 1-year decomposed root litter, to mineral SOM and (2) how these changes are altered by anthropogenic nitrogen (N) deposition by using SOM biochemical and microbiome datasets and a long-term field experiment. N deposition did not significantly alter undecomposed or 1-year decomposed root litter, but did alter decomposing microbial communities and mineral SOM biochemical composition, specifically in lignin- and lipid-derived compounds. Taken together, this restructuring of microbial communities and alteration of SOM biochemistry likely contributed to the previously observed reduction in SOM decomposition.

## 1. Introduction

Soil organic matter (SOM) constitutes the largest terrestrial organic carbon reservoir, and altered SOM decomposition dynamics under fluctuating environmental conditions could intensify or mitigate impacts of climate change. Inputs from plants, through the deposition of above- and below-ground plant residues and the release of exudates from plant roots represents the largest organic matter input to soils^1,2^. Litter chemical composition, which is often estimated in terms of its nitrogen (N) or lignin contents^3,4^, is regarded as one of the primary controls over decomposition rates and SOM accumulation and the fate of these inputs is governed by soil microbial activity^5–9^. In recent decades, there has been a shift from traditional theory of SOM chemical recalcitrance, where biochemically recalcitrant lignified material was thought to constitute the majority of SOM because of its theorized resistance to microbial decomposition^10,11^, towards a continuum model of SOM complexity that proposes microbial-derived products as the primary constituents of SOM and the potential importance of molecular diversity in enabling SOM persistence^12–15^.

Biochemical compounds that comprise litter exist across a spectrum of chemical complexity from simple to complex substrates and the relative abundance of these potential substrates for microbial metabolism can have consequences for the fate of C stored as mineral SOM^7,14,16^. When plant litter undergoes microbially mediated breakdown during decomposition, the resulting products will differ depending on the initial biochemical quality of the plant litter, as well as the composition of the microbial community decomposing that material^7,17^. The potential importance of biochemical complexity and diversity for SOM persistence lies in the different microbial requirements for metabolizing different biochemical compounds, both for compounds that are structurally dissimilar, as well as ones that are largely similar but may exhibit minor differences, such as differences in the position of functional groups (for example, *ortho-* versus *para-* benzoic acid)^15,18^. This rise in the acknowledgement and potential importance of a higher resolution understanding of SOM is a deviation from the standard practice in which compounds are typically grouped together as classes (e.g., lipids, etc.) that are assumed to behave similarly through the decomposition process. This results in individual compounds being grouped into classes that are inherently less refined. However, with the conceptual move away from stable SOM largely consisting of recalcitrant material towards a spectrum of which biochemical complexity and diversity enable SOM persistence, research has shown that certain SOM biochemical constituents are resistant to decomposition despite being in a compound class that was traditionally viewed as “labile” (ex: long-chain lipids)^19^. As such, there is increasing emphasis on the potential importance of SOM molecular complexity and diversity in driving SOM persistence^12,15,20–22^. Critically, there is a need to determine how this proposed relationship between SOM molecular diversity and persistence responds to changing climatic conditions^23^.

To better understand how anthropogenic N deposition, a pervasive driver of global environmental change, affects the cycling and storage of SOM, we conducted a 37-year field experiment to simulate future rates of anthropogenic N deposition. After 20 years of experimental N deposition inputs, we found that anthropogenic N deposition slowed the rate and extent of organic matter decomposition, resulting in the accumulation of lignified material in mineral soil derived primarily from fine root litter^24–26^. This change in SOM biochemistry in response to N deposition was associated with decreases in lignolytic fungi and positively associated with saprotrophic bacteria^24,27–29^. However, lignin, and its identifiable derivatives, often comprises ≤10% mineral SOM^19,30–33^, meaning a substantial majority of SOM biochemistry has been considered unaltered by N deposition. A higher resolution analysis of SOM chemistry (i.e., at the level of individual compounds) and the soil microbiome through decomposition and N deposition remains a critical gap in our knowledge on the persistence of SOM that built up under anthropogenic N deposition.

By using both broad (compound classes) and more refined (individual compounds) SOM biochemical and soil microbiome datasets, in the context of a long-term field study, the primary objective of this study was to assess the effect of experimental N deposition on the trajectory of organic matter decomposition and stabilization from undecayed plant root litter to mineral SOM. We assessed the effect of experimental N deposition on SOM molecular abundance, diversity, and complexity during microbial decay by establishing a decomposition gradient from undecomposed fine roots to mineral SOM. Here, we define SOM molecular diversity analogous to species richness and evenness, where molecular richness represents individual biochemical compound counts and molecular evenness describes their relative abundance. We define SOM molecular complexity as how structurally (dis)similar biochemical compounds that constitute SOM are to one another^34^. More molecularly complex SOM would thus have fewer structural similarities between its compounds. We hypothesized that experimental N deposition will elicit changes in organic matter biochemistry at the individual compound-level that are not detected when assessed at the less refined compound class-level, and these compound substitutions may have functional implications for storage of SOM in the system. We further hypothesized that changes in SOM biochemistry will be associated with changes in microbial community composition that can explain the observed biochemical changes. Finally, we hypothesize that individual compound substitutions result in altered soil organic matter biochemical diversity and molecular complexity, potentially leading to long-term changes in SOM persistence.

## 2. Methods

### 2.1. Description of study sites

The Michigan Nitrogen Deposition Gradient Study was established in 1987 to examine the effects of atmospheric N deposition on ecosystem biogeochemical processes. Spanning a 500-km distance, the study area in Michigan, USA, consists of four replicate sugar maple (*Acer saccharum*) dominated northern hardwood forest stands on sandy spodosol soil (Typic Haplorthods of the Kalkaska series). The stands contain a thick organic horizon that includes a mat of fine roots (Oe/Oa) at its boundary with the A horizon. The stands are floristically and edaphically matched and encompass the full latitudinal range of northern hardwood forests in eastern North America and the Upper Great Lakes Region^35^, allowing us to generalize our findings across this widespread ecologically and economically important ecosystem (Table S1). At each site (*n* = 4), there are six 30-m by 30-m plots, three of which received experimental N additions (30 kg NO_3_^—^N ha^−1^ yr^−1^) in the form of NaNO_3_ pellets applied annually in six equal applications throughout the growing season beginning in 1994 and three of which receive ambient atmospheric N deposition (Figure 1).

**Figure 1.**
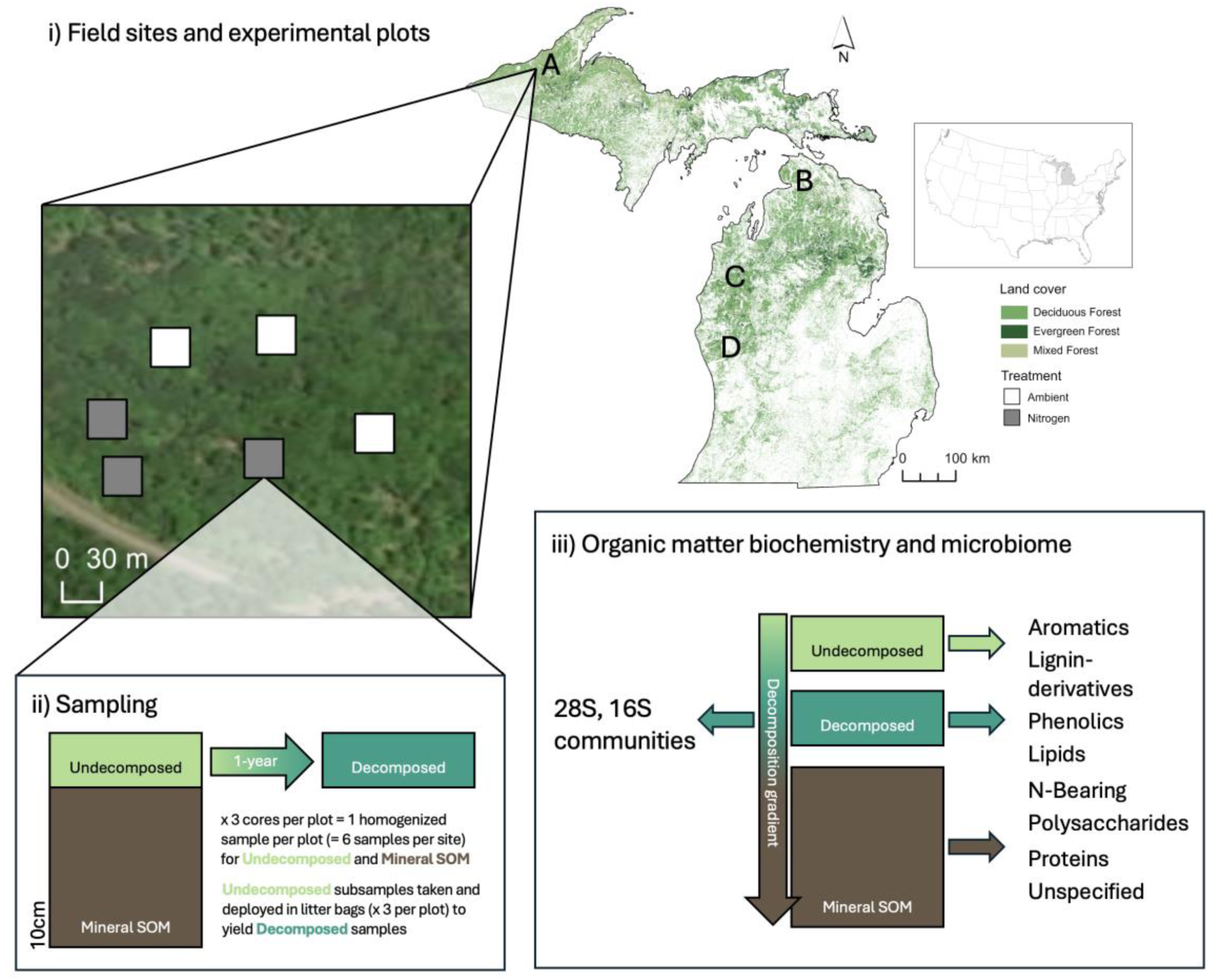
The distribution of study sites in Michigan, USA that span the geographic range of northern hardwood forests. At each site (*n* = 4), there are six 30 m x 30 m plots, three of which received annual experimental nitrogen (N) additions in the form of 30 kg NO_3_^—^N ha^−1^ yr^−1^ beginning in 1994 and three of which receive ambient atmospheric N deposition (i). Maps created using ArcGIS Pro: release 3.3.1^36^. Cores were collected (*n* = 3 per plot) to yield the undecomposed and mineral SOM samples. Subsets of undecomposed material were deployed in litter bags and left to decompose for 1-year to yield the decomposed samples (ii). The biochemistry of all three sample types was assessed using pyrolysis-GC/MS and bacterial and fungal community composition was determined on the decomposed samples (iii).

### 2.2. Field-based experiment, sampling, and sample processing

This experiment was designed to produce samples that constitute a gradient of fine root decomposition, spanning undecomposed fine roots, 1-year decomposed fine roots, and more highly decomposed mineral soil organic matter. To do this, samples were collected in September and October 2013 by collecting soil cores to a depth of 10cm (5-cm-diameter), which included both Oe/Oa and A horizons^24,37^. Cores were transported on ice to the University of Michigan and stored at −20°C. Cores were thawed and passed through a 2mm sieve, and first- through third-order fine roots were collected and pooled at the plot level^38,39^. Roots were rinsed to remove mineral soil and dried at 60°C for 24 hours. The outcome of this processing yielded the “undecomposed” and “mineral SOM” sample fractions (Figure 1). For this purpose, roots from fresh Oe/Oa horizon samples are considered undecomposed in that relative to the other sample types in this experiment, they have undergone decomposition to a lesser degree as indicated by a light coloration and lack of observable decay. Organic material in the mineral soil represents organic matter that has undergone the greatest degree of decomposition and exists as a relatively stable form of SOM compared to organic horizon material^40^.

To obtain the intermediate, “decomposed” samples, a subset of the undecomposed fine roots from each plot were sterilized and placed in mesh litter bags, deployed in the field, and collected after 12 months. Prior to field incubation, fine roots were sterilized to remove soil microbes using ethylene oxide^41^, a process that minimally alters organic matter biochemistry^42^, as compared to other methods such as autoclaving^43^. Sterilized fine roots (∼2g dry weight) were placed into 15×15cm mesh litter bags constructed with 300 µm polyester mesh on top and 20µm polyester mesh on bottom to allow for microfauna and fungal hyphae to enter the bags^37,44^. Litter bags were deployed in their respective plots to their original location in the soil profile at the boundary of the Oe/Oa and A horizons (*n* = 3 litter bags/plot at 6 plots/site; *n* = 72 litter bags total). Litter bags were placed in the field in June 2014 and collected in June 2015, immediately storing each bag on ice. The contents of each litter bag were homogenized by hand and composited at the plot level. The decomposed fine root samples were subsampled for biochemical analyses, dried at 60°C for 24 hours, and the remaining decomposed fine root material was stored at −80°C for microbial community analyses. Supporting justification and additional details on sample collection and processing can be found in Argiroff et al. (2019)^24^.

### 2.3. Organic matter biochemistry

Subsets of each of the three sample types (undecomposed fine roots, decomposed fine roots, and mineral soil; herein referred to as three different stages of decomposition) were ground with a ball mill and assessed for organic matter biochemistry using pyrolysis-GC/MS^45^. Samples were pyrolyzed at 600°C in quartz tubes for 20 seconds using a DS Pyroprobe 5150 pyrolyzer and analyzed using a ThermoTrace GC Ultra gas chromatograph (Thermo Fisher Scientific, Walthan, MA, USA) and ITQ 900 mass spectrometer (Thermo Fisher Scientific)^24,46^ at the University of New Hampshire. Compound peaks were identified and assigned using the Automated Mass Spectral Deconvolution and Identification Systems (AMDIS v2.65) software, the National Institute of Standards and Technology (NIST) compounds library, and published literature^13,45^. Organic matter compounds are expressed as the % relative abundance of total sample peak area and classified at the individual compound-level and grouped by compound source origin (e.g., lignin-derivatives, phenolics, aromatics, proteins, N-bearing, polysaccharides, and lipids). Critically, individual compounds observed in pyrolysis-GC/MS analysis are pyrolysis products of a source compound and are classified according to the nature of their source compound. Identifiable compounds with potentially multiple or unknown origins were classified as unspecified^13^. In total, this effort yielded 260 identifiable compounds, each assigned to one of the 8 compound source classes described above (Figure 1). In this study, we refer to analyses of individual pyrolysis-GC/MS compounds as “high-resolution” though we emphasize that, in this context, high-resolution refers strictly to the increased molecular resolution and biochemical information provided at the individual compound level as compared to at the broader compound class level. High-resolution analysis in this context is therefore not referring to the depth or analytical capabilities of pyrolysis-GC/MS itself relative to other high- and ultra-high resolution mass spectrometry techniques, such as FTICR-MS, used in elucidating soil organic matter biochemistry.

### 2.4. Fungal and bacterial community composition

Fungal and bacterial community composition was determined from the remaining subset of decomposed root litter and characterized using ribosomal DNA sequence abundances. Supporting justification and additional details on DNA isolation, PCR amplification, amplicon sequencing, and sequence quality control can be found in Argiroff et al. (2019)^24^. Briefly, total genomic DNA from three replicate subsamples taken from each litter bag was isolated using the DNeasy Plant Mini Kit (Qiagen, Germantown, MD) and checked for quality with a NanoDrop 8000 Spectrophotometer (Thermo Scientific) and gel electrophoresis. Replicate extractions from each litter bag were pooled and stored at −80°C prior to PCR amplification. PCR amplification was performed using the LROR and LR3 primers^47^ that target the D1-D2 region of the 28S rRNA gene. The V1-V3 regions of the bacterial 16S rRNA gene were targeted using the primers 27f and 519r^48^. PCR products were purified using the MinElute PCR Purification Kit (Qiagen) and the quality of the purified PCR products was assessed as described above. Sequencing was performed at the University of Michigan DNA Sequencing Core on 16 SMRT chips with a PacBio RS II system (Pacific Biosciences, Menlo Park, CA). Mean amplicon lengths were 688 bp for fungal 28S and 525 bp for bacterial 16S rRNA genes. Sequences were processed using mothur v1.40.5^49^. Fungal sequences were aligned against a 28S reference alignment from the RDP LSU training set^50^ and bacterial sequences were aligned against the 16S SILVA v132 reference alignment^51^. Sequences were clustered into operational taxonomic units (OTUs) for fungal and bacterial sequences at 99% and 97% sequence similarity, respectively. The most abundant sequence for each OTU was used as the representative for that OTU, and taxonomic assignments were made using the RDP classifier with the LSU training set v11 for fungi^52^ and the SILVA v132 reference alignment with the naïve Bayesian classifier^53^ in mothur for bacteria. Raw sequences are available in GenBank under the accession numbers SRR8591550 (16S) and SRR8591551 (28S).

### 2.5. Statistical Analyses

To assess the effect of experimental N deposition on the trajectory of organic matter biochemistry during decay, we assessed compositional changes elicited by the treatment in a three-step approach at two increasing levels of biochemical resolution. At the least resolved level, herein referred to as the compound class-level, we used two-way analysis of variance (ANOVA) to test the effect of experimental N deposition, site, and their interaction on log_2_-transformed compound class (*n* = 8) abundances for each of the three decomposition stages. At the higher resolution level of analysis, herein referred to as the individual compound-level, we performed PERMANOVA on Euclidean distance matrices of log_2_-transformed relative abundances of individual compounds at the three decomposition stages testing for the effect of decomposition stage, treatment, site, and their interactions on organic matter biochemistry. In the case of a significant PERMANOVA outcome, we performed PERMDISP^54^ to determine whether this effect was driven by differences in composition, heterogenous variance (i.e., dispersion), or both. The third step in our approach, at the individual compound-level, we additionally performed differential abundance analysis (Wald test)^55,56^ to test for the effect of N deposition on individual compound abundances in decomposition stages with a significant PERMANOVA treatment effect.

To assess how N deposition altered microbial community composition, we tested the effects of experimental N deposition, site, and their interaction on bacterial, fungal, and saprotroph community composition (as defined using FUNGuild in R)^57^ using two-way PERMANOVA and Bray-Curtis dissimilarity matrices calculated from Hellinger-transformed bacterial phylum, fungal genus, and saprotrophic family abundances. We performed PERMDISP to evaluate whether experimental N deposition (i.e., treatment) effects detected by PERMANOVA were driven by true differences in community composition. To determine the taxa that most differed in relative abundance across experimental N deposition, differential abundance analysis was performed on the bacterial and fungal communities using DESeq2 with *P*-value adjustment for multiple comparisons using the false discovery rate method^58,59^. To assess how N deposition altered associations between microbial communities and organic matter biochemistry, we performed Mantel tests on log_2_-transformed Euclidean distances of organic matter biochemistry and Hellinger-transformed Bray-Curtis distances of bacterial, fungal, and saprotroph communities. To determine whether changes in organic matter biochemistry were associated with changes in microbial community composition elicited by experimental N deposition, distance-based redundancy analysis (db-RDA) was performed.

We measured organic matter molecular diversity of each sample (*n* = 24 per decomposition stage; 72 total) by calculating molecular species (i.e., individual compounds) richness and evenness using the vegan package (v2.6.4)^60^ in R and performed two-way ANOVA to test the effect of decomposition stage, N deposition, and their interaction on each molecular diversity metric. Molecular richness represents individual compound counts and molecular evenness was determined by calculating Pielou’s Eveness Index^61^, which measures how equally individual compound abundances are distributed within a sample. When applicable, pairwise comparisons were made using Tukey’s honest significant difference test (HSD) to control for multiple comparisons. In contrast to molecular diversity, which assess the number of individual compounds present (i.e., richness) and their distributions within a sample (i.e., evenness), we define molecular complexity as a measure of how complex each individual compounds’ physical molecular structure is. To determine this, we calculated molecular complexity through two approaches. First, using the chemodiv package (v0.3.0)^34^ in R, we used molecular fingerprints (PubChem Fingerprint) to quantify compound dissimilarities based on the structural properties of the molecules^34^. The fingerprints were then used to calculate Jaccard dissimilarities between compounds as a measure of their structural dissimilarity^34^. We created molecular structural (dis)similarity networks to visually represent changes in organic matter molecular complexity through decomposition and between the ambient and experimental N deposition treatment. Using this method of visualization, more complex SOM would thus have fewer structural similarities between compounds and depict a less connected network. In addition to the molecular structural (dis)similarity networks, we computed molecular complexity values for each individual compound using the Bertz/Hendrickson/Ihlenfeldt formula^62,63^ computed by Cactvs 3.4.8.18 (PubChem release 2021.10.14)^64^ and performed ANOVA to test the effect of N deposition on molecular complexity. These values are estimates of how complicated an individual compound’s molecular structure is in terms of both the elements contained and structural symmetry and provide statistical significance to support network visualizations.

Due to the broad geographic expanse of our long-term field experiment and inherent heterogeneity of the soil environment, we accepted statistical significance at ⍺ = 0.1^24,27,29^. All data processing, statistical analyses, and visualizations were performed and produced in R (v4.2.2)^65^ and RStudio (version 2022.12.0+353)^66^, relying extensively on R packages such as vegan (v2.6.4)^60^, easyanova (v11.0)^67^, and the tidyverse (v1.3.1)^68^ packages.

## 3. Results

### 3.1 Compound class biochemistry

At the less-refined compound class-level, experimental N deposition significantly increased the relative abundance of lignin-derived compounds in the mineral SOM (+53%, *P* = 0.09; Figure S1; Table S2) across all sites. However, the experimental N deposition did not affect the abundance of any other compound class at any stage of decomposition (Table S3 and S4).

### 3.2 Individual compound biochemistry

When assessing the effect of experimental N deposition on the higher resolution, individual compound relative abundances across the three decomposition stages, the composition of individual compounds differed dependent on decomposition stage, site, and the N deposition treatment (decomposition stage × site × treatment; F = 1.29, *P* = 0.05; Figure 2; Table S5; Figure S2). Pairwise PERMANOVA at each decomposition stage revealed a significant effect of experimental N deposition on mineral SOM biochemistry (F = 4.26, *P* = 0.004; Figure 2; Figure S2), and no effect on the organic matter biochemistry at the undecomposed (F = 0.18, *P* = 0.93) or 1-yr decomposed (F = 0.20, *P* = 0.92) stages of decomposition (Table S5; Figure S2).

**Figure 2.**
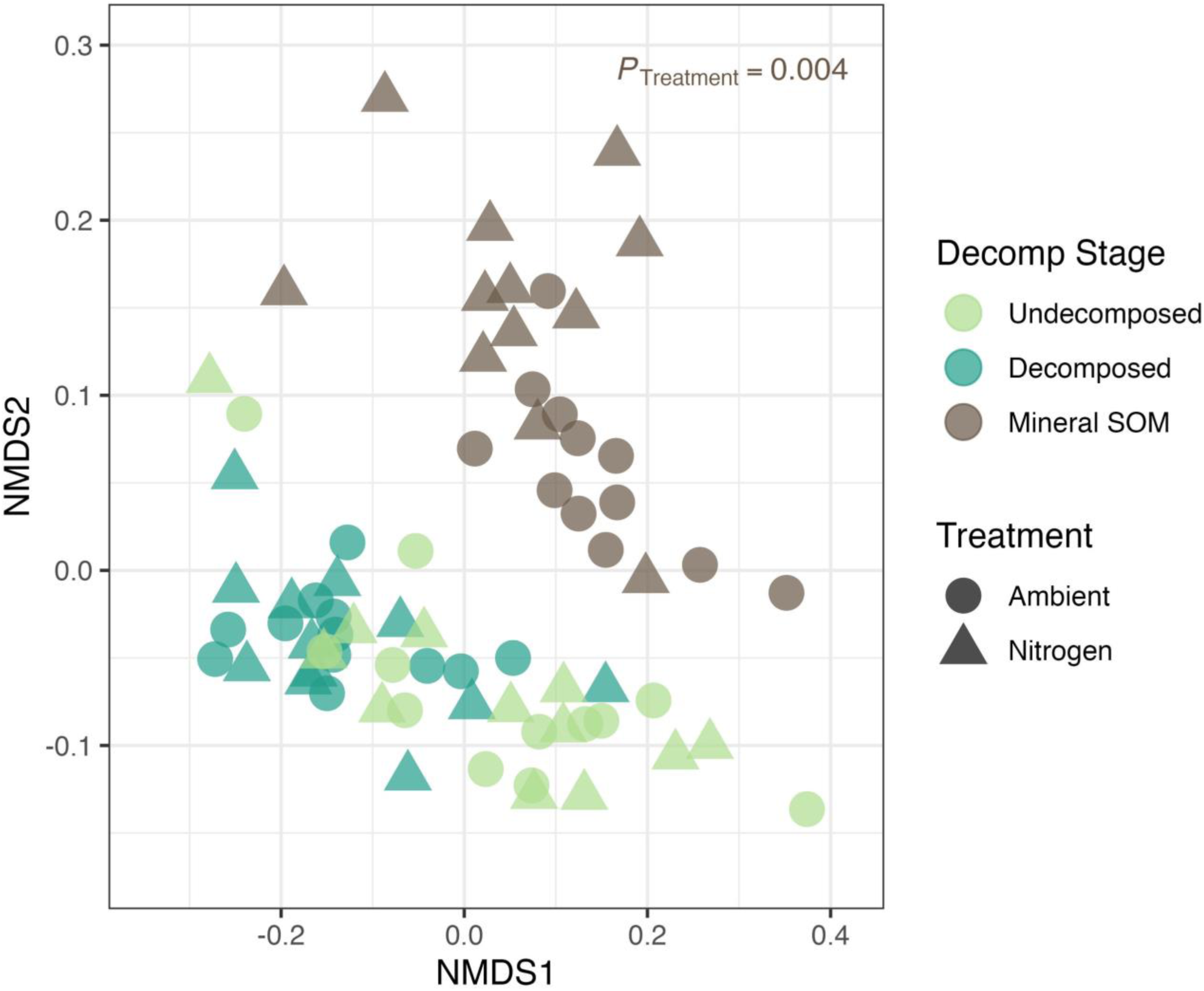
First two axes of non-metric multidimensional scaling (NMDS) ordination of Euclidean distances of log_2_ transformed py-GC/MS biochemical composition of individual samples (*n* = 72, *k* = 3, stress = 0.10). Data points represent the biochemical profile of each sample (*n* = 6 samples per site) and are colored by decomposition stage. The shape of the point represents if it received the ambient or the 30 kg NO_3_^—^N ha^−1^ yr^−1^ nitrogen deposition treatment.

Permutation tests of homogeneity of multivariate dispersion on the organic matter biochemistry at each of the three decomposition stages were nonsignificant (*P* > 0.1; Table S6), indicating that PERMANOVA detected differences in mineral SOM biochemical composition at the individual compound-level between the ambient and experimental N deposition treatment, rather than detecting changes in variability among samples (i.e., multivariate dispersion). Differential abundance tests of individual compound abundances in mineral SOM revealed that experimental N deposition led to the differential abundance of 19 out of 216 total compounds detected in the mineral SOM (Figure 3). Of these differentially abundant compounds, 7 (3.2% of the total number of compounds) were significantly more abundant in the N deposition treatment and 12 compounds (5.6% of total) were significantly more abundant in the ambient treatment (all *P* ≤ 0.1; Table S7). Compounds more abundant under experimental N deposition compared to ambient conditions included: phenol, 3-methyl- (aromatic combustion product), anthracene (aromatic), 4-guaiacylpropane (lignin), benzeneacetic acid, 4-hydroxy-3-methoxy- (lignin), benzene 1-methoxy-4-methyl- (lignin), n-decane (lipid), and 3-pyridinol (N-bearing).

**Figure 3.**
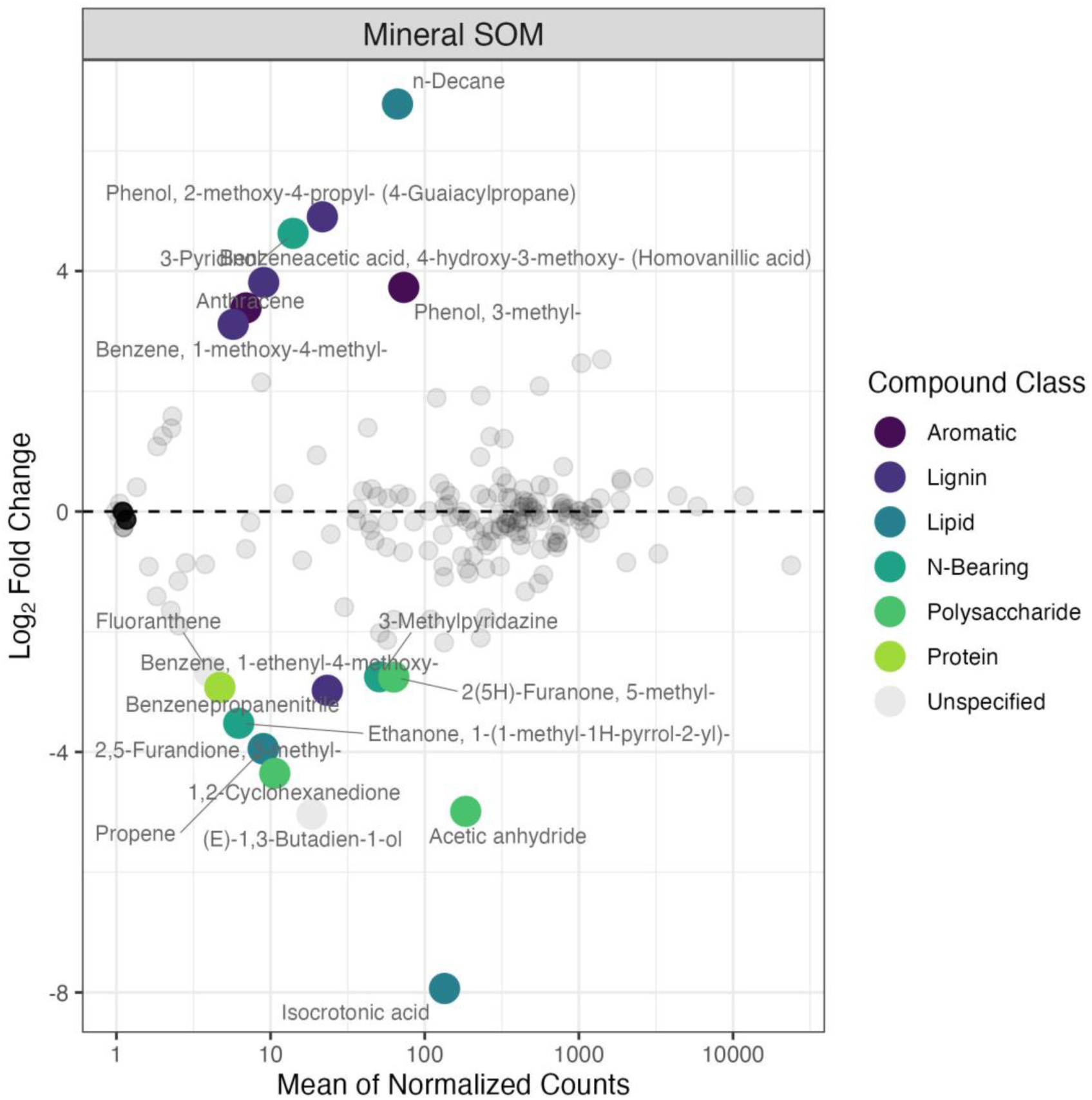
MA (Minus vs. Average) plot of mineral SOM py-GC/MS log_2_ fold changes of individual compound abundances between the ambient and nitrogen deposition treatments over the mean of normalized counts, calculated as the average normalized compound abundances taken across all samples. Compounds with a *P*_adj_ < 0.1 are colored by their compound source class. Colored compounds above the 0 line (represented by the black dashed line) indicate significantly higher abundance in the nitrogen deposition treatment, as compared to the ambient treatment, and vice versa for below the 0 line. See Table S7 for significant compound test statistics.

Compounds more abundant in the ambient treatment included: benzene 1-ethenyl-4-methoxy-(lignin), propene (lipid), isocrotonic acid (lipid), 3-methylpyridazine (N-bearing), ethanone, 1-(1-methyl-1H-pyrrol-2-yl)- (N-bearing), 2,5-furandione, 3-methyl- (N-bearing), 2(5H)-furanone, 5-methyl- (polysaccharide), 1,2-cyclohexanedione (polysaccharide), acetic anhydride (polysaccharide), benzenepropanenitrile (protein), fluoranthene (unspecified), and (E)-1,3-butadien-1-ol (unspecified).

### 3.3 Microbial community composition

Experimental N deposition significantly altered bacterial (F = 3.20, *P* = 0.009), fungal (F = 1.90, *P* = 0.017), and fungal saprotrophic community composition (F = 2.77, *P* = 0.006; Table S8; Table S9). Permutation tests of homogeneity of multivariate dispersion on each community were nonsignificant (*P* > 0.1; Table S10), indicating that PERMANOVA detected differences in microbial community composition between the ambient and experimental N deposition treatment for bacterial, fungal, and saprotrophic communities, rather than detecting changes in variability among samples (i.e., multivariate dispersion). Specifically, using the Log_2_ fold change approach in DeSeq2, the abundances of 28 bacterial OTUs (1.6% of total OTUs) were significantly higher under experimental N deposition as compared to the ambient treatment (Figure S3, Table S11). These OTUs included members of the phylum Bacteroidetes (39%), Proteobacteria (29%), Actinobacteria (14%), Patescibacteria (11%), and Armatimonadetes (7%). Under ambient conditions, 13 bacterial OTUs (0.73% of total OTUs) were present in significantly greater abundance. The phyla included Bacteroidetes (54%), Proteobacteria (38%), and Acidobacteria (8%). The abundances of 14 fungal OTUs (2.6% of total OTUs) were significantly higher under experimental N deposition as compared to the ambient treatment (Figure S3, Table S12). These OTUs included members of the fungal classes Tremellomycetes (42%), Eurotiomycetes (17%), Leotiomycetes (17%), Agaricomycetes (8%), Dothideomycetes (8%), and Sordariomycetes (8%). Under ambient conditions, 2 fungal OTUs (0.43% of total OTUs) were present in significantly greater abundance as compared to the N deposition treatment. The fungal classes include Agaricomycetes (50%) and Sordariomycetes (50%). However, the composition of bacterial (F = 1.70; P = 0.06) and fungal communities (F = 2.88; *P* < 0.001) differed among sites (Table S8; Table S9) and the effect of experimental N deposition on bacterial and fungal communities was not uniform across sites (site × treatment *P* ≤ 0.1; Table S8). Specifically, experimental N deposition did not elicit significant differences in bacterial community composition at site B (F = 2.65; *P* = 0.20) or fungal community composition at site D (F = 1.00; *P* = 0.30; Table S9).

### 3.4 Microbial community associations with SOM biochemistry

Fungal and bacterial communities were differentially associated with organic matter biochemistry at the three decomposition stages in the ambient and N deposition treatments (Figure S4). In the ambient treatment, bacterial community composition was significantly correlated with organic matter biochemistry in the undecomposed root litter (R = 0.29, *P* = 0.07) and weakly associated in the mineral SOM (R = 0.26, *P* = 0.12) but exhibited no relationship with the organic matter biochemistry in the 1-yr decomposed root litter (Table S13). Under experimental N deposition, bacterial community composition and organic matter biochemistry were not correlated at any of the three decomposition stages. In the ambient treatment, fungal community composition was also not correlated with the 1-yr decomposed root litter (Table S13) and was significantly correlated with the organic matter biochemistry in the mineral SOM (R = 0.20, *P* = 0.09). Under experimental N deposition, fungal community composition was significantly correlated with the 1-yr decomposed root litter biochemistry (R = 0.23, *P* = 0.04) and marginally associated in the mineral SOM (R = 0.17, *P* = 0.11) but exhibited no relationship with the organic matter biochemistry in the undecomposed root litter (Figure S4; Table S13). In the ambient treatment, saprotroph community composition was marginally associated with organic matter biochemistry in the undecomposed root litter (R = 0.18, *P* = 0.15) and significantly correlated with the mineral SOM (R = 0.30, *P* = 0.06) but exhibited no relationship with the organic matter biochemistry in the 1-yr decomposed root litter (Figure S4; Table S13). Under experimental N deposition, saprotroph community composition and organic matter biochemistry were not correlated at any of the three decomposition stages.

To directly link changes in organic matter biochemistry with changes in microbial community composition elicited by experimental N deposition, we fit vectors for mineral SOM individual compound abundances to bacterial (R^2^ = 0.27; *P* = 0.001), fungal (R^2^ = 0.10; *P* = 0.09), and saprotrophic (R^2^ = 0.27; *P* = 0.001) distance-based redundancy analysis (db-RDA) ordinations (Figure 4). Mineral SOM individual compound abundances were selected given the observed differences in organic matter biochemistry in the mineral SOM in response to experimental N deposition (Figure 2; Table S5; Figure 3; Table S7). The shift in bacterial community composition driven by experimental N deposition (F = 3.20, *P* = 0.009) was associated with shifts in the abundance of 28 compounds in mineral SOM (Figure 4; Table S14). The three most represented compound classes include lignin (*n* = 10; 35.7%), unspecified (*n* = 7; 25.0%), and lipids (*n* = 3; 10.7%). The longest vectors include Pyrazolo[5,1-c][1,2,4]benzotriazin-8-ol (N-bearing; vector length = 0.698), Phenol, 2,6-dimethoxy-4-(2-propenyl)- (Methoxyeugenol) (lignin; length = 0.649), Benzene, 1,4-dimethoxy- (phenol; length = 0.604), and Benzeneacetic acid, 4-hydroxy-3-methoxy- (Homovanillic acid) (lignin; length = 0.599). The shift in fungal community composition driven by experimental N deposition (F = 1.90, *P* = 0.017) was associated with shifts in the abundance of 42 compounds in mineral SOM (Figure 4; Table S15). The three most represented compound classes include lignin (*n* = 10; 23.8%), unspecified (*n* = 9; 21.4%), and lipids (*n* = 8; 19.1%). The longest vectors include Phenol, 2,6-dimethoxy-4-(2-propenyl)- (Methoxyeugenol) (lignin; length = 1.370), Ethanone, 1-(3,4-dimethoxyphenyl)- (lignin; length = 1.268), Benzoic acid, 4-hydroxy-3,5-dimethoxy-, hydrazide (unspecified; length = 1.262), and Ethanone, 1-(4-hydroxy-3,5-dimethoxyphenyl)- (Acetosyringone) (lignin; length = 1.186). The shift in saprotrophic fungal community composition under experimental N deposition (F = 2.77, *P* = 0.006) was associated with shifts in the abundance of 21 compounds in mineral SOM (Figure 4; Table S16). The three most represented compound classes include lignin (*n* = 7; 33.3%), unspecified (*n* = 7; 33.3%), and N-bearing (*n* = 3; 14.3%). The longest vectors include Trimethylphenol (unspecified; length = 1.370), Ethanone, 1-(4-hydroxy-3,5-dimethoxyphenyl)- (Acetosyringone) (lignin; length = 1.360), n-Decane (lipid; 1.350), and Benzoic acid, 4-hydroxy-3,5-dimethoxy-, hydrazide (unspecified; 1.350).

**Figure 4.**
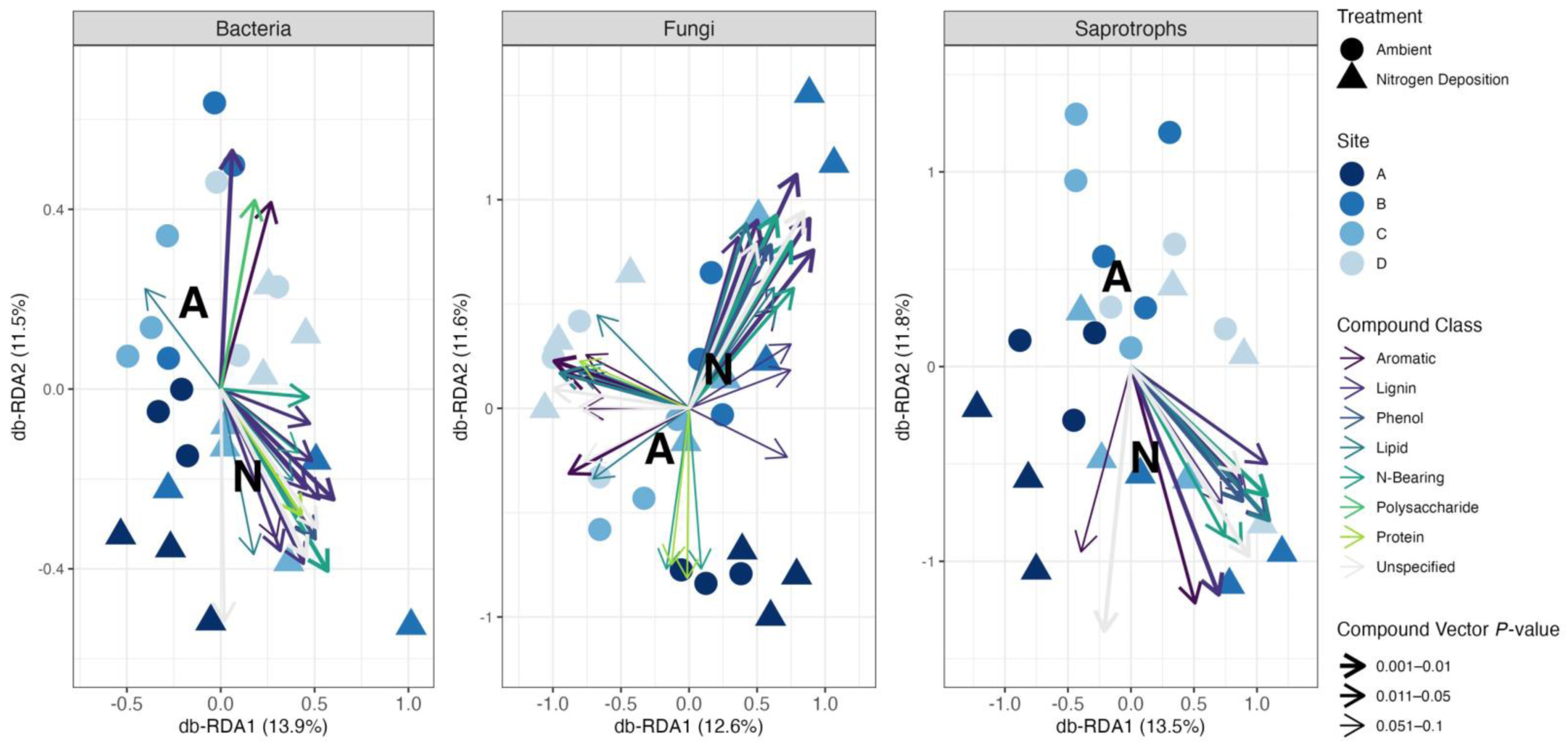
Biplots of distance-based redundancy analysis (db-RDA) ordinations of bacterial phyla, fungal genera, and saprotrophic families (points) and SOM compounds (vectors). The db-RDA was performed on Bray-Curtis dissimilarity matrices calculated from Hellinger-transformed taxon abundances. Ordinations were constrained by experimental N deposition and site, which together accounted for 31.8% of variation in Bray-Curtis dissimilarity for bacteria, 38.1% for fungi, and 33.3% for saprotrophs. Points represent individual samples (*n* = 6 samples per site) and are colored by site. Point shape indicates nitrogen treatment (ambient or 30 kg NO_3_^—^N ha^−1^ yr^−1^ deposition). Treatment centroids are shown in black as “A” (ambient) or “N” (nitrogen deposition). Vectors represent mineral SOM compounds fitted to the ordination. Compounds with goodness of fit *P*-values ≤ 0.1 are displayed and are colored by their source compound class.

### 3.5 Biochemical diversity and complexity

Experimental N deposition did not affect molecular species richness (F = 0, *P* = 0.97) or evenness (F = 0.39, *P* = 0.53) at any of the three decomposition stages (Figure 5; Table S17). Given the observed treatment differences in biochemical composition in the mineral SOM only, we assessed the effect of experimental N deposition on molecular complexity at only the mineral SOM decomposition stage. To determine how experimental N deposition affects molecular complexity (i.e., how structurally similar compounds that constitute SOM are to one another), compound co-occurrence and structural dissimilarity matrices were generated for mineral SOM biochemistry in both treatments. Under ambient conditions in the mineral SOM, molecular network analysis revealed 197 nodes, representing compounds, and 2450 edges, representing the structural similarities present between those compounds (Figure 6). Molecular network analysis of the mineral SOM biochemistry in experimental N deposition revealed it did not significantly alter the number of compounds present (*n*_nodes_ = 187) nor the structural similarities between compounds (*n*_edges_ = 2502; Figure 6). Molecular complexity values calculated using the Bertz/Hendrickson/Ihlenfeldt formula affirmed that the N deposition treatment did not significantly alter individual compound molecular complexity in the mineral SOM (F = 0, *P* = 0.997; Figure S5).

**Figure 5.**
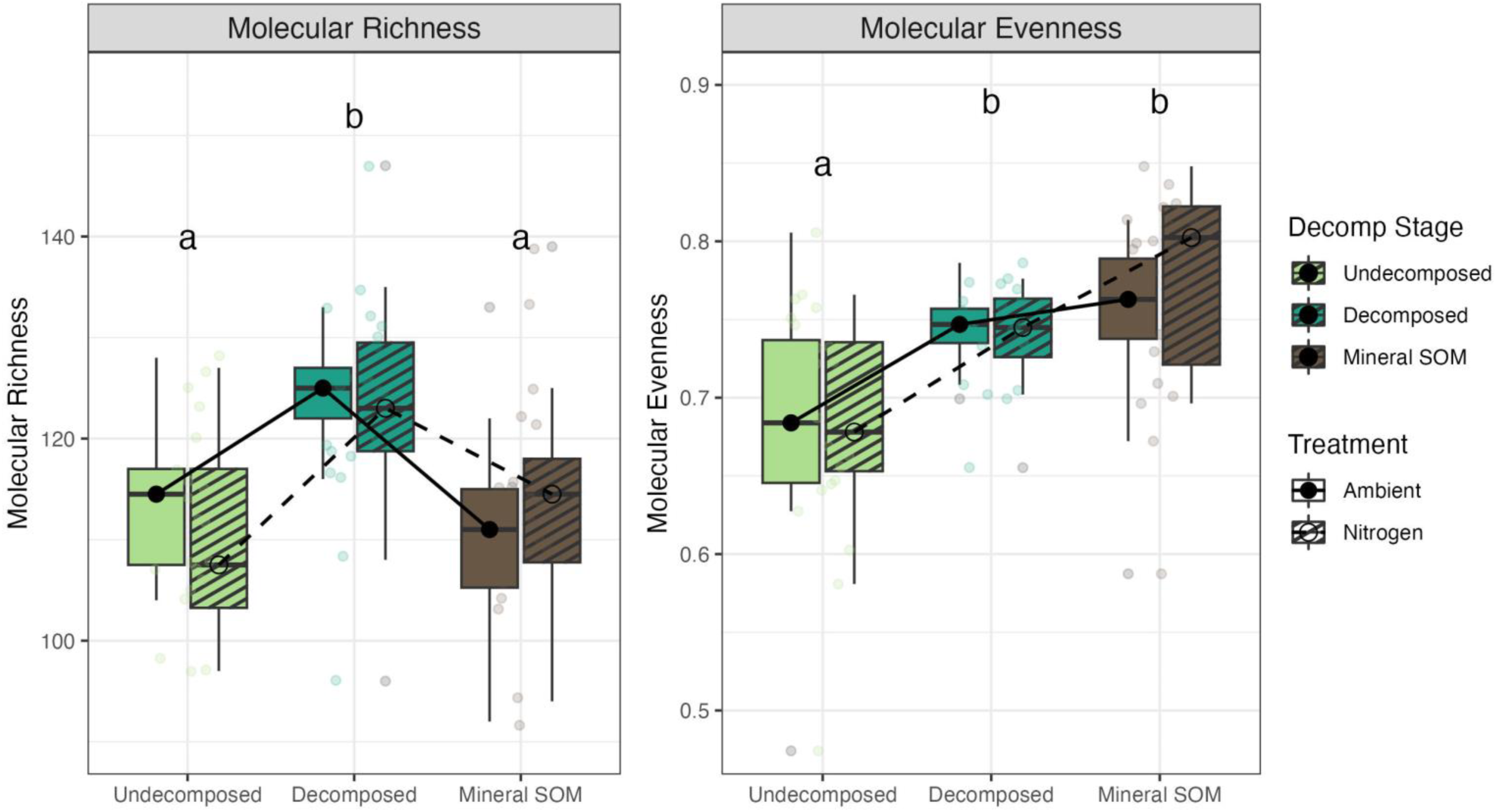
The effect of the nitrogen deposition treatment and decomposition stage on molecular richness (left) and evenness (right). Within each figure, ambient values are displayed on the left (solid) and nitrogen deposition treatment values are displayed on the right (dashed) for each decomposition stage (*n* = 3). Individual sample values (*n* = 72) are displayed. Two-way ANOVA was used to test the effect of Decomposition Stage, Treatment, and Decomposition Stage x Treatment interaction. The effect of Decomposition Stage was significant for both richness (F_decomp_ = 9.49, *P*<0.001) and evenness (F_decomp_ = 12.22, *P*<0.001). Decomposition Stage pairwise comparisons (Tukey’s HSD) are displayed as letters (see Supplementary Table S18).

**Figure 6.**
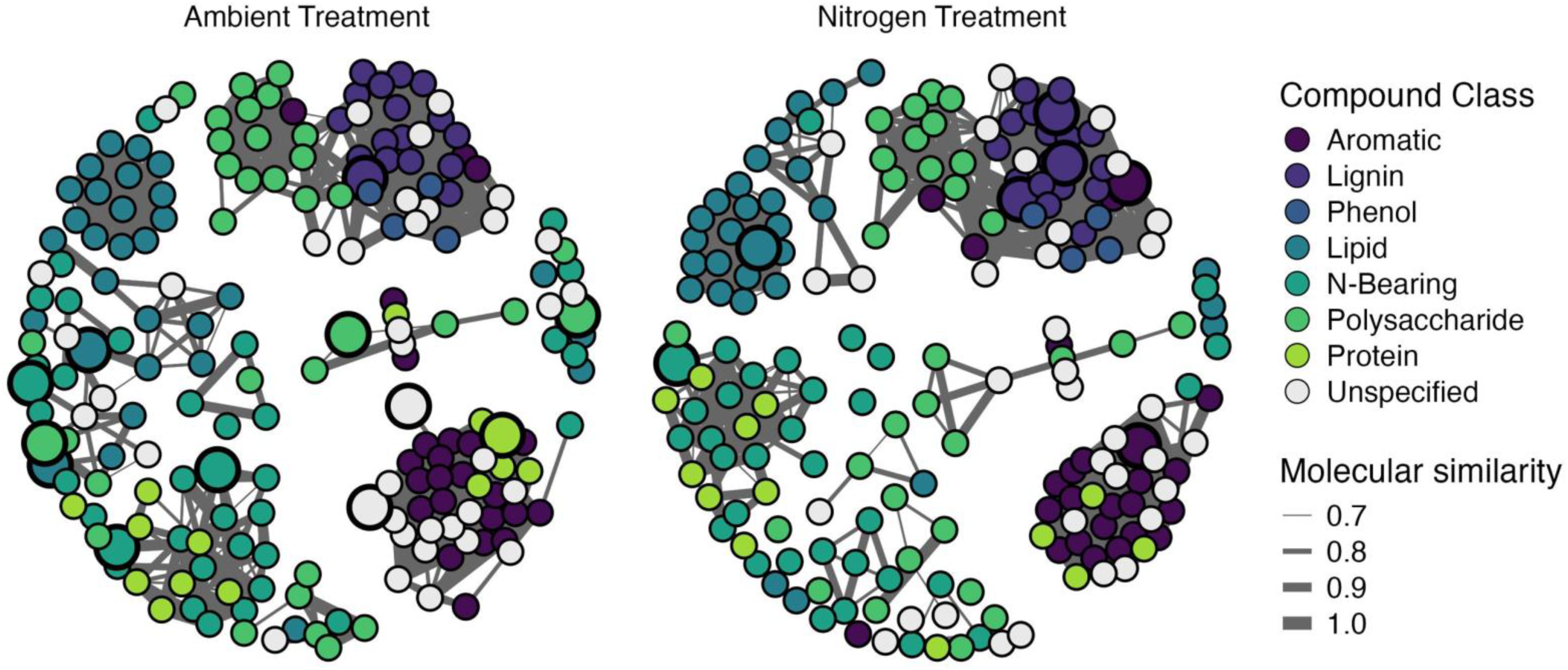
Molecular networks showing relationships between the biochemical compounds present in the mineral SOM under ambient (left) and nitrogen deposition (right) conditions. Nodes represent individual molecules (ambient: *n* = 197; nitrogen treatment: *n* = 189) and edges represent the quantity of interactions and relationships between those molecules (ambient: *n* = 2450; nitrogen treatment: *n* = 2502). Thickness of the edges was determined by the molecular similarity with a cut off value of 0.7. Nodes are colored by the compound class to which that individual molecule belongs. Enlarged nodes represent the molecules in greater abundance in each respective treatment derived from differential abundance analysis (Figure 3; Table S7).

Decomposition stage had a significant effect on molecular richness (F = 9.49, *P*<0.001) and evenness (F = 12.22, *P*<0.001; Figure 5; Table S17). HSD-adjusted pairwise comparisons revealed significantly higher molecular richness at the 1-yr decomposed stage compared to the undecomposed (post hoc *P* = 0.001) and mineral SOM (post hoc *P*<0.001; Figure 5; Table S18). Molecular evenness was significantly higher in the 1-yr decomposed (post hoc *P =* 0.002) and mineral SOM (post hoc *P*<0.001) stages of decomposition as compared to the undecomposed root litter (Figure 5; Table S18). There was no difference in molecular evenness between the 1-yr decomposed and mineral SOM decomposition stages (post hoc *P* = 0.39; Figure 5; Table S18). In the undecomposed root litter, molecular network analysis revealed 209 nodes and 2095 edges, while the 1-yr decomposed root litter network had 225 nodes and 3372 edges, and the mineral SOM network had 216 nodes and 3022 edges (Figure S6). Molecular complexity values calculated using the Bertz/Hendrickson/Ihlenfeldt formula affirmed that decomposition stage did not significantly alter individual compound molecular complexity at any point (F = 0.01, *P* = 0.99, Figure S7).

## 4. Discussion

High resolution molecular analyses reveal N deposition elicits changes to SOM biochemistry that were previously undetectable, supporting our first hypothesis. It is common to assess SOM biochemistry by grouping individual biochemical compounds into broader classes, such as lipids or lignin-derived compounds, as a means to simplify biochemical heterogeneity and diversity by grouping compounds that are suspected to behave similarly based on the relative similarity of their structure or other physical properties or factors^12,13,24,69,70^. In doing so, the abundances of individual compounds within a compound class are combined, leading to a loss in the potential resolution of SOM biochemical information, and potentially making impacts of environmental change on SOM biochemical constituents undetectable. As the resolution at which we can examine SOM biochemistry has increased with technological advances, so has the importance of identifying and elucidating the complexity within these traditional compound classes. While it has been common to assume that individual compounds within a class of compounds behave uniformly, there is increasing evidence for contrasting and compounding origins within a compound class and the different potential pathways of mineralization will have implications for the formation and persistence of mineral SOM^2,7,19,20^. As such, the implementation of more refined analytical analyses (i.e., high[er] resolution) that aid in elucidating these molecular-level differences can increase understanding of mechanisms governing SOM persistence; a necessary step toward increasing organic matter in soil as a viable climate change mitigation strategy^15,20^.

To demonstrate this assertion, previous research efforts in this long-term field experiment have detected increases in lignin-derived compounds in mineral SOM under experimental N deposition, with no detectable changes to other components of mineral SOM biochemistry^24,25^. However, high resolution analysis used in this study revealed numerous individual compounds changed in mineral SOM within multiple additional compound classes in response to experimental N deposition that may have consequences for the processes of organic matter decay and soil C storage (Figure 3; Table S7). For example, the abundance of lipids in mineral SOM exhibited no effect of experimental N deposition at the compound class-level (Figure S1; Table S2). However, at the individual compound-level, in the ambient treatment, two molecules that are produced from the combustion of lipids (propene and isocrotonic acid) were 15-times and over 200-times more abundant, respectively, compared to the experimental N deposition treatment (Figure 3; Table S7). Whereas under experimental N deposition, one lipid molecule (n-decane) was over 100-times more abundant compared to in the ambient treatment (Figure 3; Table S7) and was one of the most significant compounds associated with the shift in saprotrophic community composition driven by experimental N deposition (Figure 4; Table S16). This reorganization of individual compounds within compound classes is undetectable by less refined analysis, ultimately leading to no observable effect of experimental N deposition on the abundance of lipids as a class of compounds (Figure S1; Table S2), but is critical for understanding the fate of these substrates in SOM. Lipids are a highly heterogenous class of compounds with significant input from both plants and soil microbes^71,72^. Though there is rising evidence that stable mineral SOM is largely microbial-derived, it remains unclear and highly context dependent how much is plant-derived^13,19,20,73–76^. Thus, it is likely that observed changes in the microbial community can account for a portion of the variance observed in SOM lipid biochemistry, supporting our second hypothesis. Microbial taxa that are relatively more abundant under experimental N deposition and exhibit fast growing (i.e., copiotrophic) traits may contribute to increased lipid signatures in mineral SOM through enhanced turnover, resulting in necromass production. In this study, members of the phyla Bacteroidetes and Acidobacteria, and the genus *Paraburkholderia* were in significantly greater abundance under experimental N deposition compared to the ambient treatment (Figure S3), all of which harbor taxa that exhibit copiotrophic characteristics^77,78^. However, estimations of microbial necromass C in forest soils suggest that a substantial portion (∼70%) of C in persistent SOM could be plant-derived^79,80^, and that the contribution of plant-derived lipids to this C pool can be greater than that of lignin^20^. All three of the differentially abundant lipid molecules across the ambient and experimental N deposition treatments are involved in both plant and microbial metabolic processes^81–83^. As such, more detailed knowledge of the specific metabolic reactions, which plants or microbes are producing them, and how these compounds end up in mineral SOM is necessary to understand the longevity of these substrates in the system^84^.

Proteins and other N-containing compounds exhibited similar trends at these differing analytical scales. Despite no statistical difference in the abundance of proteins at the compound class-level, at the individual compound-level one protein molecule (benzenepropanenitrile) was almost 8-times more abundant in the ambient treatment compared under experimental N deposition (Figure 3; Table S7). The hydrolysis of benzenepropanenitrile (i.e., 3-phenylpropionitrile) is catalyzed by the nitrilase enzyme (EC 3.5.5.1)^85–88^, yielding a carboxylic acid and ammonium (NH ^+^). The gene which encodes the enzyme that ultimately carries out this reaction is found in the genomes of two soil bacteria, one of which being *Paraburkholderia phymatum*^86,89–92^. In this study, *Paraburkholderia spp.* (*n* = 3; Table S11) were in significantly greater abundance under experimental N deposition compared to the ambient treatment (Figure S3). *Paraburkholderia spp*. are known decomposers in forests, particularly sugar maple forests like the ones in this study^93^. Thus, it is plausible that increased enzymatic activity produced by *Paraburkholderia spp.* under experimental N deposition catalyzed this reaction, leading to significantly lower benzenepropanenitrile abundance in the N deposition treatment. These findings suggest that changes to microbial community composition in response to environmental stimuli may have functional consequences and elicit changes to the biochemical composition of organic matter that eventually becomes stable mineral SOM.

In addition to the novel findings detailed above, high resolution analysis of SOM biochemistry revealed additional compositional insight into lignin dynamics in response to the experimental N deposition that may underlie resistance to microbial decomposition, further supporting our first hypothesis. Previous work in this long-term experiment showed that anthropogenic N deposition slowed the rate and extent of organic matter decomposition, ultimately resulting in the accumulation of lignified material in mineral SOM under experimental N deposition (+53%, *P* = 0.09; Figure S1; Table S2)^24–26,29^. However, at the individual compound-level, changes in the composition of lignin compounds in mineral SOM were detected under both ambient and experimental N deposition conditions. In the ambient treatment, one lignin-derived molecule (benzene, 1-ethenyl-4-methoxy-) was almost 8-times more abundant compared to under experimental N deposition (Figure 3; Table S7). Whereas under experimental N deposition, three different lignin-derived molecules (benzene, 1-methoxy-4-methyl-, homovanillic acid, and 4-guaiacylpropane) were 8-times, 14-times, and 29-times more abundant than in the ambient treatment, respectively, ultimately leading to the overall increase observed at the compound class-level (Figure 3; Figure S1; Table S7).

At the relatively less-refined compound class-level, it is not possible to detect this reorganization of different lignin molecules, as observed here, that ultimately affects the interpretability of the stability of the compound class as a whole. Lignin is a complex macromolecule that is highly variable in terms of its primary structure. Which lignin monomers and functional groups are present, in what quantities, and what linkages connect them affect the decomposability of lignin polymers by soil microbes^94–97^. For example, lignin polymers containing higher quantities of monomers of the guaiacyl (G) subunit are more resistant to decomposition, resulting in significantly lower mass loss, than lignin polymers composed of other subunits^16,96^. The 29-times greater abundance of 4-guaiacylpropane and 14-times greater abundance of homovanillic acid, a G subunit decomposition product^98^, under experimental N deposition suggests that the SOM that has accumulated under anthropogenic N deposition is particularly compositionally resistant to microbial decomposition in terms of its lignin content. Thus, it is plausible that in this system, differences in the quality of lignin, in addition to the increases in lignin quantity^24^, may be a factor contributing to the increased SOM observed under high N deposition conditions. This possibility has been suggested from earlier modeling studies in this system^26^, and this is the first experimental observation to support this modeled result.

Differences in the composition of decomposing microbial communities observed under experimental N deposition support the claim that lignin in mineral SOM under increased anthropogenic N deposition is particularly resistant to decomposition or is simply not being metabolized by the microbial community in this treatment, further supporting our second hypothesis. While fungi are widely considered as primary lignin decomposers, evidence has shown that bacteria may play a non-inconsequential role in lignin decay^99,100^. Of the bacterial OTUs in higher abundance under N deposition conditions, just under half (43%) belong to the phyla Proteobacteria and Actinobacteria (Figure S3; Table S11). This finding aligns with previous work in this system that observed an increase in the abundance of bacteria associated with lignin decay under experimental N deposition^27,28^. The majority of lignin-degrading bacteria identified to date represent three bacterial classes, all of which are within the Proteobacteria and Actinobacteria phyla: Actinomycetes, *α*-Proteobacteria, and *γ*-Proteobacteria^101–103^. Such bacteria capable of lignin degradation do so in an incomplete manner, such that smaller aromatic lignin-derived compounds are produced^104–106^. For example, bacterial species belonging to the class *γ*-Proteobacteria produce the intermediate phenolic compound homovanillic acid during plant litter and lignin decomposition^107^. In this study, homovanillic acid was one of the most significant compounds associated with the shift in bacterial community composition driven by experimental N deposition (Figure 4; Table S14) and was in significantly higher abundance in the N deposition treatment compared to the ambient treatment (Figure 3; Table S7).

Additionally, benzoic acid, 4-hydroxy-3,5-dimethoxy-, hydrazide, a decomposition derivative of syringic acid, which is a key intermediate of bacterial catabolism of lignin compounds^108^, was one of the most significant compounds associated with the shifts in fungal and saprotrophic community composition driven by experimental N deposition (Figure 4; Table S15; Table S16). Taken together, it is plausible that the accumulation of decomposition resistant lignin material under experimental N deposition was due, in part, to the incomplete bacterial metabolism of lignin and the inability to break down these compounds specifically.

Further, both bacterial and saprotrophic fungal communities exhibit strong relationships with the biochemistry of the organic matter in the undecomposed litter and mineral SOM under ambient conditions (Figure S4; Table S13). Yet, under experimental N deposition these relationships are lost. The expression of microbial genes and the subsequent production of extracellular enzymes that govern the breakdown of SOM are highly sensitive to N additions^29,109–112^. Under high N conditions, genes responsible for the production of extracellular enzymes that break down complex macromolecules, such as lignin, are downregulated^29^.

Additional studies have observed a shift in microbial community composition under N deposition conditions towards a community whose primary niche is the decay of simpler compounds, such as cellulose^109,113,114^. Taken together, it is plausible that when N is readily available, microbes do not need to expend the energy producing extracellular enzymes to break down organic material and this decreased activity may result in a shift in the community composition. The lack of microbial activity observed in previous studies, in consideration with SOM biochemical relationships to microbial taxa observed here, may explain why we observed a decoupling of SOM biochemistry and microbial community composition under experimental N deposition.

Despite the above-described differences in individual compound biochemical composition being distinct under N deposition conditions in the most highly decomposed mineral SOM (Figure 3; Table S7), the biochemical diversity and molecular complexity of mineral SOM as a whole was not altered by N deposition (Figure 5; Figure 6), contradicting our third hypothesis. However, SOM biochemical diversity (i.e., molecular richness and evenness) did change throughout decomposition (Figure 5). Both molecular richness and evenness were higher after one year of decomposition compared to initial undecomposed inputs (Figure 5; Table S17; Table S18). This is likely the result of the production of microbially-derived compounds during the initial phases of root decay. When microbes break down plant litter, new byproducts are produced^7,13,115^, which likely explains the observed increases in molecular richness, i.e., the number of compounds present, after one year of decomposition. Molecular richness of mineral SOM returning to the same levels as initial inputs may indicate the complete decay of more accessible components of litter or that the microbially processed C substrates that were produced during one year of decomposition were similarly consumed, returning to the same number of compounds as initial inputs, though the composition of those compounds are distinct (Figure 2; Figure 5). Despite differences in biochemical diversity through decomposition, we did not detect differences in the molecular complexity of SOM between the undecomposed root litter, 1-year decomposed root litter, and mineral SOM (Figure S6; Figure S7). Ultimately these findings indicate that although the composition of the individual compounds changed during decay (Figure 2; Table S5), and were ultimately distinct under N deposition conditions in the most highly decomposed mineral SOM (Figure 2; Figure 3; Table S7), the molecular complexity (i.e., structural [dis]similarity) of the compounds that constitute SOM at any given stage of decomposition did not (Figure S4). This finding differs from both what has been proposed conceptually about organic matter complexity underlying SOM persistence^15^ and from recent studies that have found the opposite: that persistent SOM is less complex than its initial inputs^12,21,116^. In similar studies, SOM persistence was estimated across wider ecosystem ranges (e.g., across soil depths or elevational gradients that span different ecosystem types^12,21,116^) than in this study, which utilized a long-term experiment in sugar maple dominated forests. Thus, in this system, the lack of relationship between SOM molecular complexity and persistence may be attributable to relatively similar site-specific edaphic factors and vegetation composition, which may not be variable enough to detect differences in SOM decay using this method that detects large-scale molecular changes (e.g., terpenes vs. benzenoids). Taken together, our findings suggest that SOM biochemical diversity and molecular complexity are not drivers of SOM persistence in this system.

Collectively, our results demonstrate that N deposition altered the abundance of specific compounds in mineral SOM, a reorganization of SOM biochemistry that is not detectable when assessing changes at the less-refined compound class-level, as well as decoupled the relationship between microbial composition and SOM biochemistry. Whether these observed changes have functional implications for the long-term storage of SOM is unknown and warrants further investigation. Though these molecular-level differences in SOM biochemistry under anthropogenic N deposition may appear subtle, the effect that these differences have on the C stored in the system long-term may be disproportionately large. For example, in this system, upon termination of the experimental N deposition, a globally ecologically relevant environmental change in response to Clean Air Act Amendments, microbial activity significantly increased and the C that built up under experimental N deposition was lost from the system within 5-years of treatment termination^40^. Studies have shown that molecular-level changes in SOM biochemistry, in response to various environmental change conditions, impact the capacity of temperate forests to sequester C^117^. A microbial mechanism may underlie this possibility as differences in the structural properties of the molecules in greater abundance under anthropogenic N deposition may have implications for how and by whom these products are transformed, what products are produced, and, ultimately, the decomposability of their byproducts^83,118,119^. Taken together, it is plausible that the SOM that accumulated under experimental N deposition was molecularly distinct as a result of functional changes in the microbial community elicited by the N deposition treatment^24,29^. Thus, upon termination of experimental N deposition, the accumulated, molecularly distinct SOM may have been readily decomposed by the active microbial community upon return to low (i.e., ambient) N conditions. While empirically untested, this response is plausible due to the relative increase in lignin-like compounds, yet observed decrease in the expression of genes that encode lignolytic enzymes^29^, as previously observed under experimental N deposition. As such, lignolytic fungal gene expression may rapidly increase in response to reduced N deposition, supporting a rapid decay of the relatively lignin-rich SOM that accumulated during the active experimental N deposition treatment in this long-term study. Future work should explore the relative decomposition potential of different lignin- and lipid-related compounds that were differentially abundant under ambient or the experimental N deposition treatment to deepen our understanding of the potential ecosystem-level consequences of the reorganization of SOM chemistry observed here.

## Acknowledgements

We thank Annalisa Stevenson for her help in ArcGIS. We also wish to thank Thea Whitman (and lab), Erika Marín-Spiotta (and lab), and the Department of Soil Science graduate students for their thoughtful feedback on this work.

## Author Contributions Statement

D.R.Z. designed the study. R.A.U. and W.A.A. performed field and laboratory analyses. A.S.G. performed biochemical analyses. B.E.P. analyzed the data with the help of G.A.C and Z.B.F. B.E.P wrote the first draft of the manuscript. D.R.Z., W.A.A., G.A.C., A.S.G., and Z.B.F. provided significant feedback on subsequent versions and analyses.

