## Supplemental Tables and Figures for "Anthropogenic nitrogen deposition restructures decomposing microbial communities, altering SOM molecular composition, but not molecular complexity or diversity"

[1]Department of Soil and Environmental Sciences
University of Wisconsin, Madison, WI, USA 53706
[2]University of Michigan Biological Station
9133 Biological Rd, Pellston, MI, USA 49769
[3]School for Environment and Sustainability
[4]Ecology and Evolutionary Biology
University of Michigan, Ann Arbor, MI, USA 48109
[5]Department of Natural Resources and the Environment
University of New Hampshire, Durham, NH, USA 03824

**Table S1.** Site, climate, vegetation, and soil characteristics of the four northern hardwood forest stands in Michigan, USA (see Figure 1). The sites exist across a natural north-south climatic gradient, yet are similar in age, plant composition, and soil development.

| Characteristic | Site A | Site B | Site C | Site D |
|---|---|---|---|---|
| *Location* | | | | |
| Latitude (N) | 46°52' | 45°33' | 44°23' | 43°40' |
| Longitude (W) | 88°53' | 84°52' | 85°50' | 86°09' |
| *Climate* | | | | |
| Mean annual temperature (°C) | 4.7 | 6.0 | 6.9 | 7.6 |
| Mean annual precipitation (mm) | 873 | 871 | 888 | 812 |
| Ambient wet + dry total N deposition (g $m^{-2}$ $yr^{-1}$) | 0.68 | 0.91 | 1.17 | 1.18 |
| Growing season length (days) | 134 | 150 | 154 | 157 |
| *Vegetation* | | | | |
| Overstory biomass (Mg $ha^{-1}$) | 261 | 261 | 274 | 234 |
| *Acer saccharum* biomass (Mg $ha^{-1}$) | 237 | 224 | 216 | 201 |
| *Soil* | | | | |
| Soil texture (0-10cm) (%sand-%silt-%clay) | 75:22:3 | 89:9:2 | 89:9:2 | 87:10:3 |

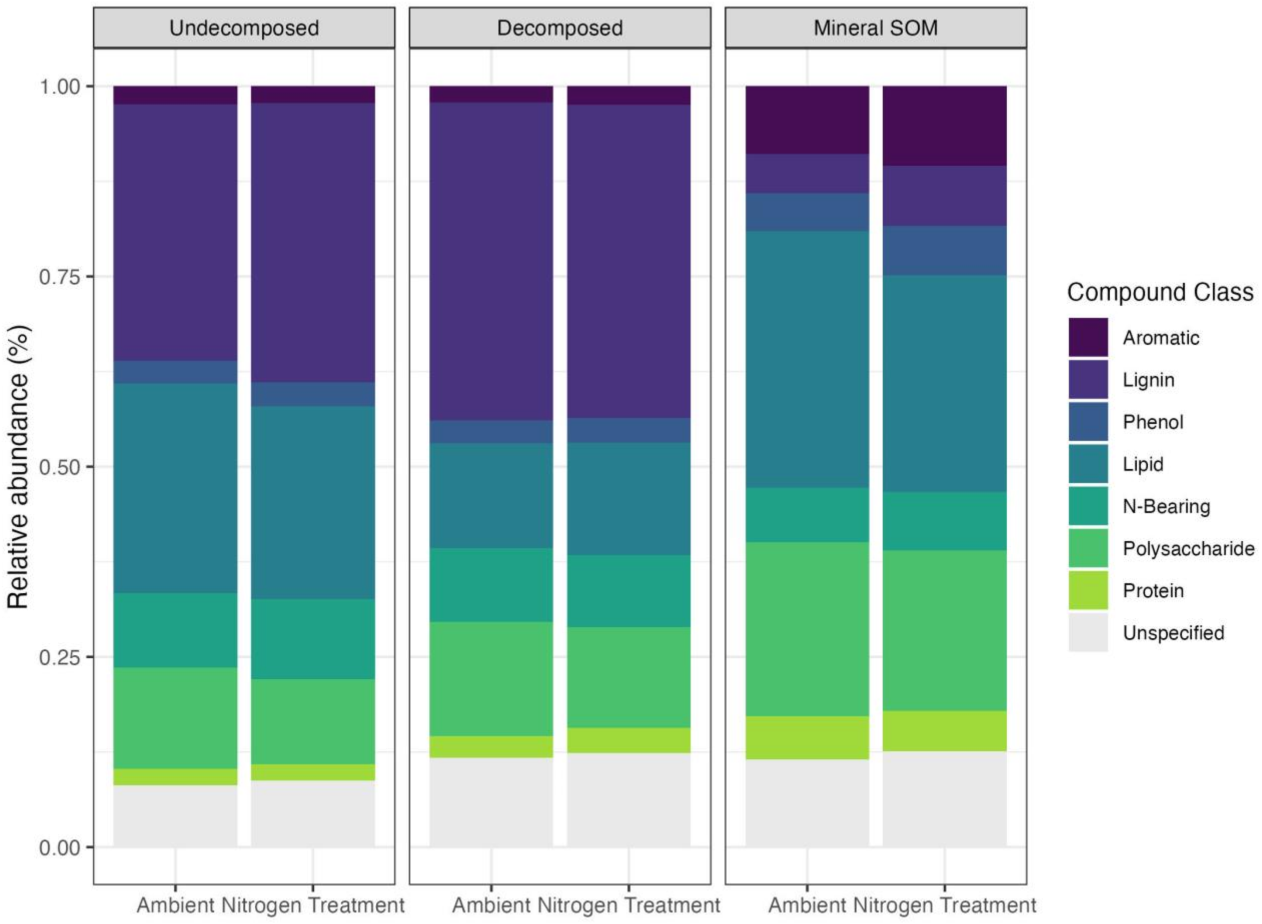


**Figure S1.** Stacked abundance chart of the relative abundances of the 8 compound classes across the three decomposition stages from least (left, undecomposed) to most (right, mineral SOM) decomposed under ambient and simulated nitrogen deposition conditions. Across the three decomposition stages, the nitrogen deposition treatment only significantly affected the relative abundance of lignin in the mineral SOM (+53% in nitrogen treatment, $P = 0.09$).

**Table S2.** Two-way ANOVA results for the effect of Site, Treatment, and Site x Treatment interaction on $\log_2$-transformed relative abundances of the 8 compounds classes in mineral Soil Organic Matter (SOM). Significant P-values ($P \leq 0.1$) are presented in bold.

| Compound class | Factor | Numerator df | Denominator df | F-value | P-value |
|---|---|---|---|---|---|
| Aromatic | Site | 3 | 20 | 2.5041 | **0.0962** |
| | Treatment | 1 | 22 | 1.4847 | 0.2407 |
| | Site x Treatment | 3 | 20 | 0.5880 | 0.6316 |
| Lignin | Site | 3 | 20 | 6.3903 | **0.0047** |
| | Treatment | 1 | 22 | 3.2040 | **0.0924** |
| | Site x Treatment | 3 | 20 | 0.4467 | 0.7230 |
| Lipid | Site | 3 | 20 | 1.2469 | 0.3257 |
| | Treatment | 1 | 22 | 1.5861 | 0.2259 |
| | Site x Treatment | 3 | 20 | 0.1884 | 0.9027 |
| N-Bearing | Site | 3 | 20 | 7.3233 | **0.0026** |
| | Treatment | 1 | 22 | 0.8199 | 0.3786 |
| | Site x Treatment | 3 | 20 | 2.9805 | **0.0626** |
| Phenol | Site | 3 | 20 | 0.6791 | 0.5775 |
| | Treatment | 1 | 22 | 2.9435 | 0.1055 |
| | Site x Treatment | 3 | 20 | 0.4292 | 0.7349 |
| Polysaccharide | Site | 3 | 20 | 0.5502 | 0.6552 |
| | Treatment | 1 | 22 | 0.4042 | 0.5339 |
| | Site x Treatment | 3 | 20 | 4.5208 | **0.0177** |
| Protein | Site | 3 | 20 | 1.7234 | 0.2024 |
| | Treatment | 1 | 22 | 0.5559 | 0.4667 |
| | Site x Treatment | 3 | 20 | 0.3023 | 0.8233 |
| Unspecified | Site | 3 | 20 | 1.1552 | 0.3574 |
| | Treatment | 1 | 22 | 0.4160 | 0.5281 |
| | Site x Treatment | 3 | 20 | 0.8866 | 0.4691 |

**Table S3.** Two-way ANOVA results for the effect of Site, Treatment, and Site x Treatment interaction on $log_2$-transformed relative abundances of the 8 compounds classes in undecomposed fine roots. Significant P-values ($P \leq 0.1$) are presented in bold.

| Compound class | Factor | Numerator df | Denominator df | F-value | P-value |
|---|---|---|---|---|---|
| Aromatic | Site | 3 | 20 | 0.9337 | 0.4473 |
| | Treatment | 1 | 22 | 0.2609 | 0.6165 |
| | Site x Treatment | 3 | 20 | 2.1511 | 0.1338 |
| Lignin | Site | 3 | 20 | 0.0870 | 0.9662 |
| | Treatment | 1 | 22 | 0.4924 | 0.4929 |
| | Site x Treatment | 3 | 20 | 1.2213 | 0.3343 |
| Lipid | Site | 3 | 20 | 1.1512 | 0.3588 |
| | Treatment | 1 | 22 | 0.1842 | 0.6735 |
| | Site x Treatment | 3 | 20 | 1.1179 | 0.3711 |
| N-Bearing | Site | 3 | 20 | 2.7075 | **0.0799** |
| | Treatment | 1 | 22 | 0.4525 | 0.5108 |
| | Site x Treatment | 3 | 20 | 1.1713 | 0.3516 |
| Phenol | Site | 3 | 20 | 1.5606 | 0.2378 |
| | Treatment | 1 | 22 | 0.3278 | 0.5749 |
| | Site x Treatment | 3 | 20 | 4.5536 | **0.0173** |
| Polysaccharide | Site | 3 | 20 | 4.3391 | **0.0203** |
| | Treatment | 1 | 22 | 1.5337 | 0.2334 |
| | Site x Treatment | 3 | 20 | 3.9589 | **0.0275** |
| Protein | Site | 3 | 20 | 0.7584 | 0.5337 |
| | Treatment | 1 | 22 | 0.0003 | 0.9860 |
| | Site x Treatment | 3 | 20 | 0.5795 | 0.6369 |
| Unspecified | Site | 3 | 20 | 1.5001 | 0.2526 |
| | Treatment | 1 | 22 | 0.3898 | 0.5412 |
| | Site x Treatment | 3 | 20 | 1.3106 | 0.3054 |

**Table S4.** Two-way ANOVA results for the effect of Site, Treatment, and Site x Treatment interaction on $log_2$-transformed relative abundances of the 8 compounds classes in the 1-yr decomposed fine roots. Significant P-values ($P \leq 0.1$) are presented in bold.

| Compound class | Factor | Numerator df | Denominator df | F-value | P-value |
|---|---|---|---|---|---|
| Aromatic | Site | 3 | 20 | 3.5241 | **0.0393** |
| | Treatment | 1 | 22 | 0.7394 | 0.4026 |
| | Site x Treatment | 3 | 20 | 1.4160 | 0.2747 |
| Lignin | Site | 3 | 20 | 0.3242 | 0.8078 |
| | Treatment | 1 | 22 | 0.0587 | 0.8116 |
| | Site x Treatment | 3 | 20 | 1.4453 | 0.2668 |
| Lipid | Site | 3 | 20 | 1.1059 | 0.3757 |
| | Treatment | 1 | 22 | 0.0909 | 0.7669 |
| | Site x Treatment | 3 | 20 | 1.5248 | 0.2464 |
| N-Bearing | Site | 3 | 20 | 2.2232 | 0.1250 |
| | Treatment | 1 | 22 | 0.1254 | 0.7279 |
| | Site x Treatment | 3 | 20 | 0.2880 | 0.8334 |
| Phenol | Site | 3 | 20 | 2.2061 | 0.1270 |
| | Treatment | 1 | 22 | 0.4187 | 0.5267 |
| | Site x Treatment | 3 | 20 | 0.6662 | 0.5849 |
| Polysaccharide | Site | 3 | 20 | 2.5255 | **0.0943** |
| | Treatment | 1 | 22 | 1.3879 | 0.2560 |
| | Site x Treatment | 3 | 20 | 0.5719 | 0.6416 |
| Protein | Site | 3 | 20 | 2.4745 | **0.0988** |
| | Treatment | 1 | 22 | 2.6923 | 0.1203 |
| | Site x Treatment | 3 | 20 | 4.2287 | **0.0222** |
| Unspecified | Site | 3 | 20 | 0.1201 | 0.9469 |
| | Treatment | 1 | 22 | 0.3110 | 0.5848 |
| | Site x Treatment | 3 | 20 | 0.2234 | 0.8787 |

**Table S5.** Three-way PERMANOVA results for the effect of Decomposition Stage, Site, Treatment, and their interactions on $log_2$-transformed Euclidean distance matrices of individual compound abundances. Below the three-way PERMANOVA results are pairwise two-way PERMANOVA results for the effect of Site, Treatment, and their interaction at the three decomposition stages. Significant P-values ($P \leq 0.1$) are presented in bold.

| Decomp Stage | Factor | Df | $R^2$ | F-value | P-value |
|---|---|---|---|---|---|
| | Decomp Stage | 2 | 0.35930 | 20.707 | **0.001** |
| | Site | 3 | 0.01930 | 0.7416 | 0.645 |
| | Treatment | 1 | 0.01351 | 1.5572 | 0.193 |
| | Decomp Stage x Site | 6 | 0.06012 | 1.1548 | 0.312 |
| | Decomp Stage x Treatment | 2 | 0.02397 | 1.3816 | 0.211 |
| | Site x Treatment | 3 | 0.01440 | 0.5531 | 0.843 |
| | Decomp Stage x Site x Treatment | 6 | 0.09296 | 1.7858 | **0.048** |
| Undecomposed | Site | 3 | 0.12377 | 0.9610 | 0.444 |
| | Treatment | 1 | 0.00765 | 0.1783 | 0.934 |
| | Site x Treatment | 3 | 0.17265 | 1.4107 | 0.212 |
| Decomposed | Site | 3 | 0.13225 | 1.0294 | 0.405 |
| | Treatment | 1 | 0.00869 | 0.2028 | 0.924 |
| | Site x Treatment | 3 | 0.17384 | 1.3530 | 0.244 |
| Mineral SOM | Site | 3 | 0.11883 | 1.0912 | 0.362 |
| | Treatment | 1 | 0.15467 | 4.2612 | **0.004** |
| | Site x Treatment | 3 | 0.14573 | 1.3382 | 0.197 |

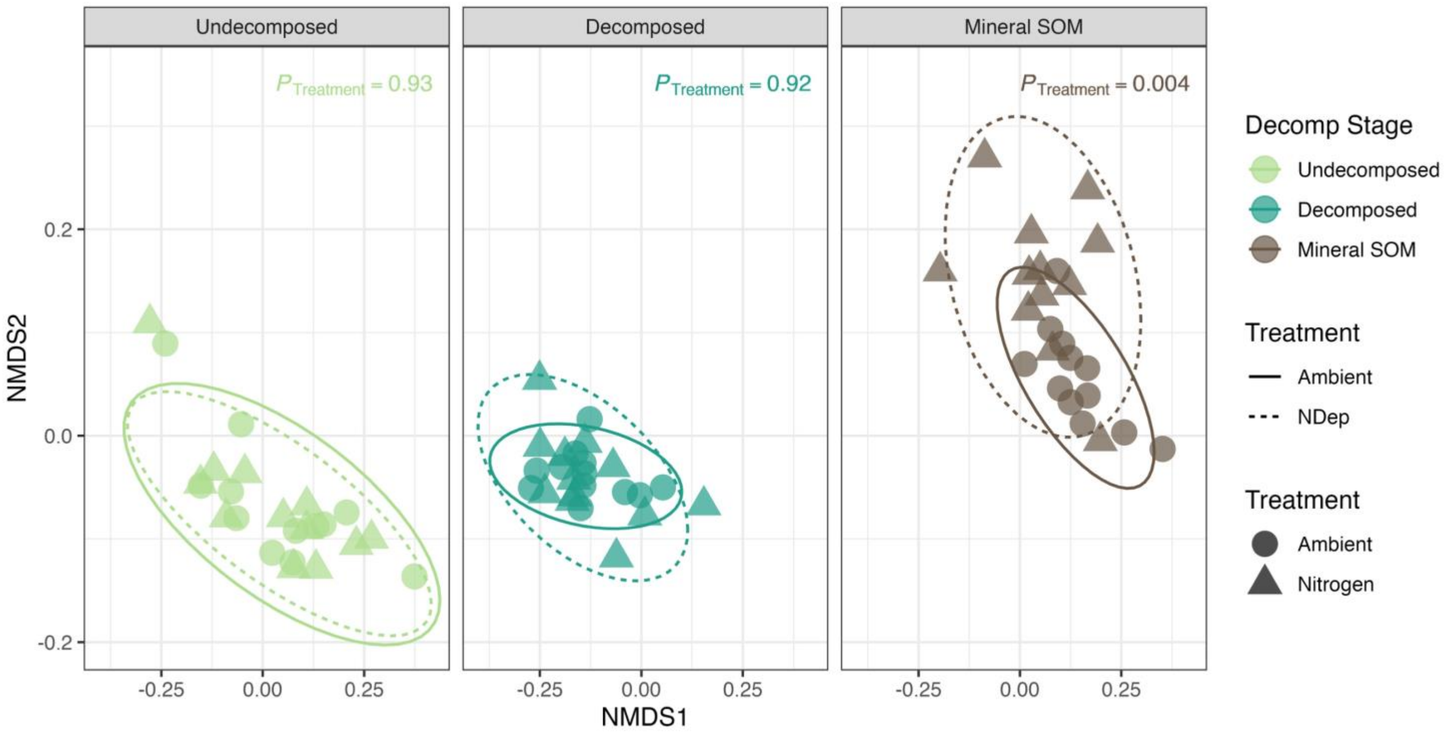


**Figure S2.** First two axes of non-metric multidimensional scaling (NMDS) ordination of Euclidean distances of $log_2$ transformed py-GC/MS biochemical composition of individual samples displayed in separate panels for each decomposition stage ($n$ = 72, $k$ = 3, stress = 0.10). Data points represent the biochemical profile of each sample ($n$ = 6 samples per site) and are colored by decomposition stage. The shape of the point represents if it received the ambient or the 30 kg $NO_3^{-}$–N $ha^{-1}$ $yr^{-1}$ nitrogen deposition treatment. Ellipses represent 95% confidence intervals calculated for each treatment (ambient = solid, N deposition = dashed) in each decomposition stage.

**Table S6.** Permutation test for homogeneity of multivariate dispersion results for the effect of Treatment on $log_2$-transformed Euclidean distance matrices of individual compound abundances at each of the three decomposition stages.

| Decomp Stage | Factor | Df | F-value | P-value |
|---|---|---|---|---|
| Undecomposed | Treatment | 1 | 0.0949 | 0.756 |
| Decomposed | Treatment | 1 | 0.2859 | 0.612 |
| Mineral SOM | Treatment | 1 | 1.7029 | 0.213 |

**Table S7.** Mineral SOM compounds with significant differential abundance (Benjamini-Hochberg $P_{\mathrm{adj}} \leq 0.1$) and $Log_2$ fold change values that are plotted in Figure 3. Treatment refers to the treatment that the compound exhibits significantly greater abundance in.

| Compound | Compound Class | Treatment | $Log_2$ Fold Change | Standard Error | Wald Statistic | BH $P_{adj}$ |
|---|---|---|---|---|---|---|
| Phenol, 3-methyl- | Aromatic | Nitrogen | 3.72780 | 1.12256 | 3.32081 | 0.01017 |
| Anthracene | Aromatic | Nitrogen | 3.38703 | 0.94245 | 3.59388 | 0.00462 |
| Phenol, 2-methoxy-4-propyl- (4-Guaiacylpropane) | Lignin | Nitrogen | 4.90243 | 0.97139 | 5.04680 | 0.00003 |
| Benzeneacetic acid, 4-hydroxy-3-methoxy- (Homovanillic acid) | Lignin | Nitrogen | 3.81029 | 0.94943 | 4.01324 | 0.00127 |
| Benzene, 1-methoxy-4-methyl- | Lignin | Nitrogen | 3.11383 | 0.90764 | 3.43070 | 0.00787 |
| Benzene, 1-ethenyl-4-methoxy- | Lignin | Ambient | -2.97359 | 1.01665 | -2.92489 | 0.03446 |
| n-Decane | Lipid | Nitrogen | 6.77821 | 1.05588 | 6.41951 | <0.0001 |
| Propene | Lipid | Ambient | -3.95030 | 1.01705 | -3.88408 | 0.00183 |
| Isocrotonic acid | Lipid | Ambient | -7.93390 | 1.07392 | -7.38778 | <0.0001 |
| 3-Pyridinol | N-Bearing | Nitrogen | 4.62658 | 1.06348 | 4.35040 | 0.00039 |
| 3-Methylpyridazine | N-Bearing | Ambient | -2.74810 | 1.00508 | -2.73422 | 0.05905 |
| Ethanone, 1-(1-methyl-1H-pyrrol-2-yl)- | N-Bearing | Ambient | -3.52273 | 0.92806 | -3.79581 | 0.00227 |
| 2,5-Furandione, 3-methyl- | N-Bearing | Ambient | -3.94198 | 1.01805 | -3.87210 | 0.00183 |
| 2(5H)-Furanone, 5-methyl- | Polysaccharide | Ambient | -2.75395 | 1.07045 | -2.57270 | 0.09029 |
| 1,2-Cyclohexanedione | Polysaccharide | Ambient | -4.35563 | 1.04552 | -4.16602 | 0.00075 |
| Acetic anhydride | Polysaccharide | Ambient | -4.98762 | 1.09738 | -4.54501 | 0.00019 |
| Benzenepropanenitrile | Protein | Ambient | -2.92576 | 0.86588 | -3.37895 | 0.00884 |
| Fluoranthene | Unspecified | Ambient | -2.66957 | 0.83642 | -3.19167 | 0.01503 |
| (E)-1,3-Butadien-1-ol | Unspecified | Ambient | -5.03650 | 1.09046 | -4.61868 | 0.00016 |

**Table S8.** Two-way PERMANOVA results for the effect of the N deposition treatment, site, and their interaction on Hellinger-transformed Bray-Curtis dissimilarity distance matrices of bacterial phylum, fungal genus, and saprotrophic family abundances. Significant P-values ($P \leq 0.1$) are presented in bold.

| Type | Factor | Df | $R^2$ | F-value | P-value |
|---|---|---|---|---|---|
| Bacteria (16S) | Site | 3 | 0.20345 | 1.7027 | **0.0601** |
| | Treatment | 1 | 0.12707 | 3.2024 | **0.0096** |
| | Site x Treatment | 7 | 0.50648 | 2.3457 | **0.0022** |
| Fungi (28S) | Site | 3 | 0.30172 | 2.8806 | **0.0001** |
| | Treatment | 1 | 0.07957 | 1.9018 | **0.0169** |
| | Site x Treatment | 7 | 0.51496 | 2.4268 | **0.0001** |
| Saprotrophs | Site | 3 | 0.23087 | 2.0012 | **0.0067** |
| | Treatment | 1 | 0.11205 | 2.7760 | **0.0068** |
| | Site x Treatment | 7 | 0.47219 | 2.0448 | **0.0001** |

**Table S9.** Pairwise PERMANOVA results where two-way PERMANOVA of microbial community composition (Table S8) revealed significant ($P \leq 0.1$) effect of site, treatment, and/or site x treatment interaction. Significant P-values ($P \leq 0.1$) are presented in bold.

| Type | Factor | Post-hoc Comparison | $R^2$ | F | P-value |
|---|---|---|---|---|---|
| Bacteria (16S) | Site | A – B | 0.500 | 2.66 | **0.0084** |
| | | A – C | 0.453 | 2.21 | **0.0023** |
| | | A – D | 0.609 | 4.16 | **0.0002** |
| | | B – C | 0.441 | 2.10 | **0.0490** |
| | | B – D | 0.424 | 1.97 | **0.0520** |
| | | C – D | 0.438 | 2.08 | **0.0083** |
| | Site x Treatment | Ambient A – Nitrogen A | 0.443 | 3.19 | **0.1000** |
| | | Ambient B – Nitrogen B | 0.398 | 2.65 | 0.2000 |
| | | Ambient C – Nitrogen C | 0.343 | 2.09 | **0.1000** |
| | | Ambient D – Nitrogen D | 0.341 | 2.07 | **0.1000** |
| Fungi (28S) | Site | A – B | 0.496 | 2.62 | **0.0001** |
| | | A – C | 0.465 | 2.32 | **0.0001** |
| | | A – D | 0.436 | 2.07 | **0.0004** |
| | | B – C | 0.493 | 2.60 | **0.0011** |
| | | B – D | 0.455 | 2.23 | **0.0002** |
| | | C – D | 0.420 | 1.93 | **0.0022** |
| | Site x Treatment | Ambient A – Nitrogen A | 0.272 | 1.49 | **0.1000** |
| | | Ambient B – Nitrogen B | 0.384 | 2.50 | **0.1000** |
| | | Ambient C – Nitrogen C | 0.360 | 2.25 | **0.1000** |
| | | Ambient D – Nitrogen D | 0.199 | 1.00 | 0.3000 |
| Saprotrophs | Site | A – B | 0.503 | 2.40 | **0.0008** |
| | | A – C | 0.394 | 1.74 | **0.0220** |
| | | A – D | 0.389 | 1.70 | **0.0440** |
| | | B – C | 0.504 | 2.71 | **0.0046** |
| | | B – D | 0.408 | 1.83 | **0.0350** |
| | | C – D | 0.355 | 1.47 | **0.0930** |
| | Site x Treatment | Ambient A – Nitrogen A | 0.194 | 0.97 | 0.6000 |
| | | Ambient B – Nitrogen B | 0.490 | 3.85 | **0.1000** |
| | | Ambient C – Nitrogen C | 0.352 | 2.17 | 0.2000 |
| | | Ambient D – Nitrogen D | 0.155 | 0.734 | 0.7000 |

**Table S10.** Permutation test for homogeneity of multivariate dispersion results for the effect of Treatment on Hellinger-transformed Bray-Curtis dissimilarity distance matrices of bacterial phylum, fungal genus, and saprotrophic family abundances.

| Type | Factor | Df | F-value | P-value |
|---|---|---|---|---|
| Bacteria (16S) | Treatment | 1 | 2.3530 | 0.1368 |
| Fungi (28S) | Treatment | 1 | 1.2156 | 0.2715 |
| Saprotrophs | Treatment | 1 | 2.6555 | 0.1172 |

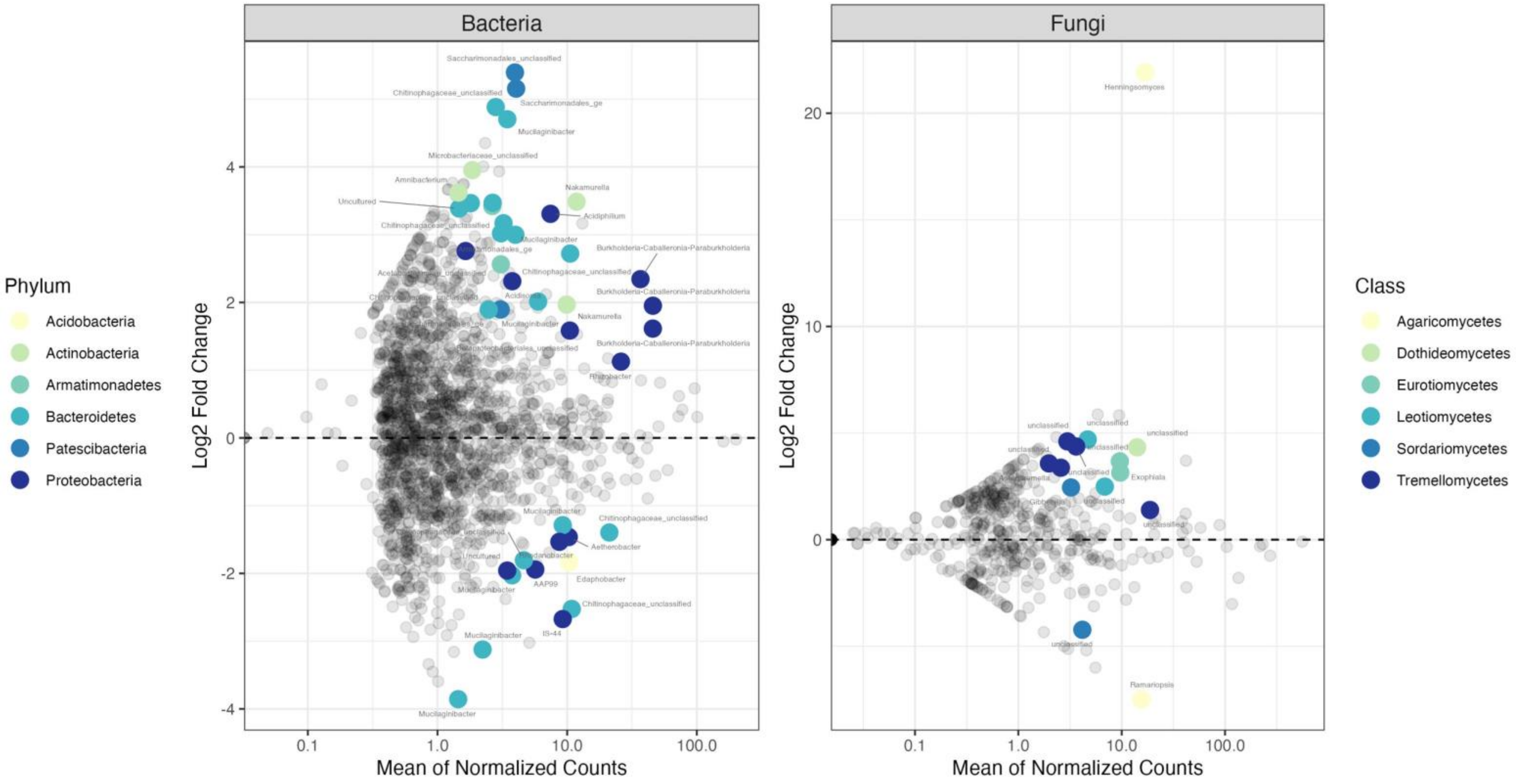


**Figure S3.** Plot of the $Log_2$ Fold Change over the Mean of Normalized Counts for the bacteria (left) and fungi (right) genera comparing species abundance between the ambient and nitrogen deposition treatments in the "decomposed" samples. Species with a $p_{adj} < 0.1$ are colored by phylum. Colored species above the 0 line indicate a significantly higher abundance in the nitrogen deposition treatment, as compared to the ambient treatment, and vice versa for below the 0 line.

**Table S11.** Bacteria with significant differential abundance (Benjamini-Hochberg $P_{\mathrm{adj}} \leq 0.1$) and $\mathrm{Log}_2$ fold change values that are plotted in Figure S3 (left). Treatment refers to the treatment that the OTU (species) exhibits significantly greater abundance in.

| Kingdom | Phylum | Class | Order | Family | Genus | Treatment | $\mathrm{Log}_2$ Fold Change | Standard Error | Wald Statistic | BH $P_{adj}$ |
|---|---|---|---|---|---|---|---|---|---|---|
| Bacteria | Patescibacteria | Saccharimonadia | Saccharimonadales | Unclassified | Unclassified | Nitrogen | 5.39412 | 1.50297 | 3.58898 | 0.01505 |
| Bacteria | Patescibacteria | Saccharimonadia | Saccharimonadales | Unclassified | Unclassified | Nitrogen | 5.15561 | 1.19097 | 4.32891 | 0.00322 |
| Bacteria | Bacteroidetes | Bacteroidia | Chitinophagales | Chitinophagaceae | Unclassified | Nitrogen | 4.88380 | 1.29715 | 3.76504 | 0.01029 |
| Bacteria | Bacteroidetes | Bacteroidia | Sphingobacteriales | Sphingobacteriaceae | Mucilaginibacter | Nitrogen | 4.70389 | 0.84858 | 5.54322 | 0.00001 |
| Bacteria | Actinobacteria | Actinobacteria | Micrococcales | Microbacteriaceae | Unclassified | Nitrogen | 3.95287 | 1.29021 | 3.06375 | 0.05308 |
| Bacteria | Actinobacteria | Actinobacteria | Micrococcales | Microbacteriaceae | Amnibacterium | Nitrogen | 3.62117 | 1.18129 | 3.06544 | 0.05308 |
| Bacteria | Actinobacteria | Actinobacteria | Frankiales | Nakamurellaceae | Nakamurella | Nitrogen | 3.48676 | 1.03957 | 3.35404 | 0.02843 |
| Bacteria | Bacteroidetes | Bacteroidia | Chitinophagales | Chitinophagaceae | Ferruginibacter | Nitrogen | 3.47086 | 1.25134 | 2.77371 | 0.09754 |
| Bacteria | Bacteroidetes | Bacteroidia | Chitinophagales | Chitinophagaceae | Unclassified | Nitrogen | 3.46616 | 1.15698 | 2.99588 | 0.06041 |
| Bacteria | Armatimonadetes | Armatimonadia | Armatimonadales | Unclassified | Unclassified | Nitrogen | 3.42300 | 0.93394 | 3.66513 | 0.01401 |
| Bacteria | Bacteroidetes | Bacteroidia | Chitinophagales | Chitinophagaceae | Uncultured | Nitrogen | 3.38620 | 0.95645 | 3.54038 | 0.01688 |
| Bacteria | Proteobacteria | Alphaproteobacteria | Acetobacterales | Acetobacteraceae | Acidiphilium | Nitrogen | 3.30733 | 0.93798 | 3.52600 | 0.01688 |
| Bacteria | Bacteroidetes | Bacteroidia | Chitinophagales | Chitinophagaceae | Unclassified | Nitrogen | 3.16901 | 1.14426 | 2.76950 | 0.09754 |
| Bacteria | Bacteroidetes | Bacteroidia | Chitinophagales | Chitinophagaceae | Unclassified | Nitrogen | 3.02005 | 1.05278 | 2.86863 | 0.08495 |
| Bacteria | Bacteroidetes | Bacteroidia | Sphingobacteriales | Sphingobacteriaceae | Mucilaginibacter | Nitrogen | 2.99831 | 1.07783 | 2.78179 | 0.09754 |
| Bacteria | Proteobacteria | Alphaproteobacteria | Acetobacterales | Acetobacteraceae | Unclassified | Nitrogen | 2.75872 | 0.95259 | 2.89603 | 0.08031 |
| Bacteria | Bacteroidetes | Bacteroidia | Chitinophagales | Chitinophagaceae | Unclassified | Nitrogen | 2.72105 | 0.90886 | 2.99392 | 0.06041 |
| Bacteria | Armatimonadetes | Armatimonadia | Armatimonadales | Unclassified | Unclassified | Nitrogen | 2.56260 | 0.74517 | 3.43895 | 0.02206 |
| Bacteria | Proteobacteria | Gammaproteobacteria | Betaproteobacteriales | Burkholderiaceae | Burkholderia-Caballeronia-Paraburkholderia | Nitrogen | 2.34356 | 0.62196 | 3.76802 | 0.01029 |
| Bacteria | Proteobacteria | Alphaproteobacteria | Acetobacterales | Acetobacteraceae | Acidisoma | Nitrogen | 2.31260 | 0.59484 | 3.88776 | 0.00860 |
| Bacteria | Bacteroidetes | Bacteroidia | Chitinophagales | Chitinophagaceae | Uncultured | Nitrogen | 3.38620 | 0.95645 | 3.54038 | 0.01688 |
| Bacteria | Proteobacteria | Alphaproteobacteria | Acetobacterales | Acetobacteraceae | Acidiphilium | Nitrogen | 3.30733 | 0.93798 | 3.52600 | 0.01688 |
| Bacteria | Bacteroidetes | Bacteroidia | Sphingobacteriales | Sphingobacteriaceae | Mucilaginibacter | Nitrogen | 2.01288 | 0.51097 | 3.93931 | 0.00794 |
| Bacteria | Actinobacteria | Actinobacteria | Frankiales | Nakamurellaceae | Nakamurella | Nitrogen | 1.96852 | 0.60164 | 3.27192 | 0.03310 |
| Bacteria | Proteobacteria | Gammaproteobacteria | Betaproteobacteriales | Burkholderiaceae | Burkholderia-Caballeronia-Paraburkholderia | Nitrogen | 1.95134 | 0.62661 | 3.11412 | 0.04825 |

| | | | | | | | | | | |
|---|---|---|---|---|---|---|---|---|---|---|
| Bacteria | Patescibacteria | Saccharimonadia | Saccharimonadales | Unclassified | Unclassified | Nitrogen | 1.89504 | 0.68840 | 2.75283 | 0.09799 |
| Bacteria | Bacteroidetes | Bacteroidia | Chitinophagales | Chitinophagaceae | Unclassified | Nitrogen | 1.89319 | 0.67628 | 2.79941 | 0.09754 |
| Bacteria | Proteobacteria | Gammaproteobacteria | Betaproteobacteriales | Burkholderiaceae | Burkholderia-Caballeronia-Paraburkholderia | Nitrogen | 1.61355 | 0.44282 | 3.64382 | 0.01405 |
| Bacteria | Proteobacteria | Gammaproteobacteria | Betaproteobacteriales | Unclassified | Unclassified | Nitrogen | 1.58444 | 0.48965 | 3.23585 | 0.03436 |
| Bacteria | Proteobacteria | Gammaproteobacteria | Betaproteobacteriales | Burkholderiaceae | Rhizobacter | Nitrogen | 1.12582 | 0.29837 | 3.77320 | 0.01029 |
| Bacteria | Bacteroidetes | Bacteroidia | Sphingobacteriales | Sphingobacteriaceae | Mucilaginibacter | Ambient | -1.28789 | 0.40941 | -3.14574 | 0.04506 |
| Bacteria | Bacteroidetes | Bacteroidia | Chitinophagales | Chitinophagaceae | Unclassified | Ambient | -1.39623 | 0.42964 | -3.24976 | 0.03415 |
| Bacteria | Proteobacteria | Deltaproteobacteria | Myxococcales | Polyangiaceae | Aetherobacter | Ambient | -1.46319 | 0.40563 | -3.60718 | 0.01504 |
| Bacteria | Proteobacteria | Gammaproteobacteria | Xanthomonadales | Rhodanobacteraceae | Rhodanobacter | Ambient | -1.53376 | 0.38863 | -3.94662 | 0.00794 |
| Bacteria | Bacteroidetes | Bacteroidia | Chitinophagales | Chitinophagaceae | Unclassified | Ambient | -1.80558 | 0.45541 | -3.96475 | 0.00794 |
| Bacteria | Acidobacteria | Acidobacteriia | Acidobacteriales | Acidobacteriaceae_ | Edaphobacter | Ambient | -1.83235 | 0.64683 | -2.83280 | 0.09228 |
| Bacteria | Proteobacteria | Gammaproteobacteria | Betaproteobacteriales | Burkholderiaceae | AAP99 | Ambient | -1.94233 | 0.45411 | -4.27721 | 0.00322 |
| Bacteria | Proteobacteria | Alphaproteobacteria | Micropepsales | Micropepsaceae | Uncultured | Ambient | -1.95804 | 0.65177 | -3.00419 | 0.06041 |
| Bacteria | Bacteroidetes | Bacteroidia | Sphingobacteriales | Sphingobacteriaceae | Mucilaginibacter | Ambient | -2.02805 | 0.60709 | -3.34058 | 0.02843 |
| Bacteria | Bacteroidetes | Bacteroidia | Chitinophagales | Chitinophagaceae | Unclassified | Ambient | -2.52434 | 0.44489 | -5.67414 | 0.00001 |
| Bacteria | Proteobacteria | Gammaproteobacteria | Betaproteobacteriales | Nitrosomonadaceae | IS-44 | Ambient | -2.67385 | 0.95610 | -2.79662 | 0.09754 |
| Bacteria | Bacteroidetes | Bacteroidia | Sphingobacteriales | Sphingobacteriaceae | Mucilaginibacter | Ambient | -3.12362 | 1.13077 | -2.76239 | 0.09754 |
| Bacteria | Bacteroidetes | Bacteroidia | Sphingobacteriales | Sphingobacteriaceae | Mucilaginibacter | Ambient | -3.85663 | 1.17895 | -3.27124 | 0.03310 |

**Table S12.** Fungi with significant differential abundance (Benjamini-Hochberg $P_{\mathrm{adj}} \leq 0.1$) and $\mathrm{Log}_2$ fold change values that are plotted in Figure S3 (right). Treatment refers to the treatment that the OTU (species) exhibits significantly greater abundance in.

| Kingdom | Phylum | Class | Order | Family | Genus | Treatment | $\mathrm{Log}_2$ Fold Change | Standard Error | Wald Statistic | BH $P_{adj}$ |
|---|---|---|---|---|---|---|---|---|---|---|
| Fungi | Basidiomycota | Agaricomycetes | Agaricales | Schizophyllaceae | Henningsomyces | Nitrogen | 21.90978 | 2.94575 | 7.43776 | <0.00001 |
| Fungi | Ascomycota | Leotiomycetes | Unclassified | Unclassified | Unclassified | Nitrogen | 4.69786 | 1.43680 | 3.26967 | 0.03446 |
| Fungi | Basidiomycota | Tremellomycetes | Tremellales | Unclassified | Unclassified | Nitrogen | 4.60797 | 1.72523 | 2.67094 | 0.09310 |
| Fungi | Basidiomycota | Tremellomycetes | Tremellales | Unclassified | Unclassified | Nitrogen | 4.36903 | 1.65903 | 2.63349 | 0.09659 |
| Fungi | Ascomycota | Dothideomycetes | Pleosporales | Unclassified | Unclassified | Nitrogen | 4.33137 | 1.34815 | 3.21282 | 0.03505 |
| Fungi | Ascomycota | Eurotiomycetes | Chaetothyriales | Herpotrichiellaceae | Unclassified | Nitrogen | 3.67041 | 1.26887 | 2.89266 | 0.06651 |
| Fungi | Basidiomycota | Tremellomycetes | Tremellales | Unclassified | Unclassified | Nitrogen | 3.57464 | 1.02756 | 3.47875 | 0.02015 |
| Fungi | Basidiomycota | Tremellomycetes | Tremellales | Tremellaceae | Asterotremella | Nitrogen | 3.37403 | 1.20852 | 2.79186 | 0.06987 |
| Fungi | Ascomycota | Eurotiomycetes | Chaetothyriales | Herpotrichiellaceae | Exophiala | Nitrogen | 3.14462 | 1.09721 | 2.86602 | 0.06651 |
| Fungi | Ascomycota | Leotiomycetes | Helotiales | Unclassified | Unclassified | Nitrogen | 2.48835 | 0.79990 | 3.11083 | 0.04264 |
| Fungi | Ascomycota | Sordariomycetes | Hypocreales | Nectriaceae | Gibberella | Nitrogen | 2.44656 | 0.87541 | 2.79476 | 0.06987 |
| Fungi | Basidiomycota | Tremellomycetes | Tremellales | Tremellaceae | Unclassified | Nitrogen | 1.39111 | 0.38410 | 3.62174 | 0.01808 |
| Fungi | Ascomycota | Sordariomycetes | Unclassified | Unclassified | Unclassified | Ambient | -4.22650 | 1.47236 | -2.87056 | 0.06651 |
| Fungi | Basidiomycota | Agaricomycetes | Agaricales | Clavariaceae | Ramariopsis | Ambient | -7.48238 | 2.08804 | -3.58345 | 0.01808 |

86

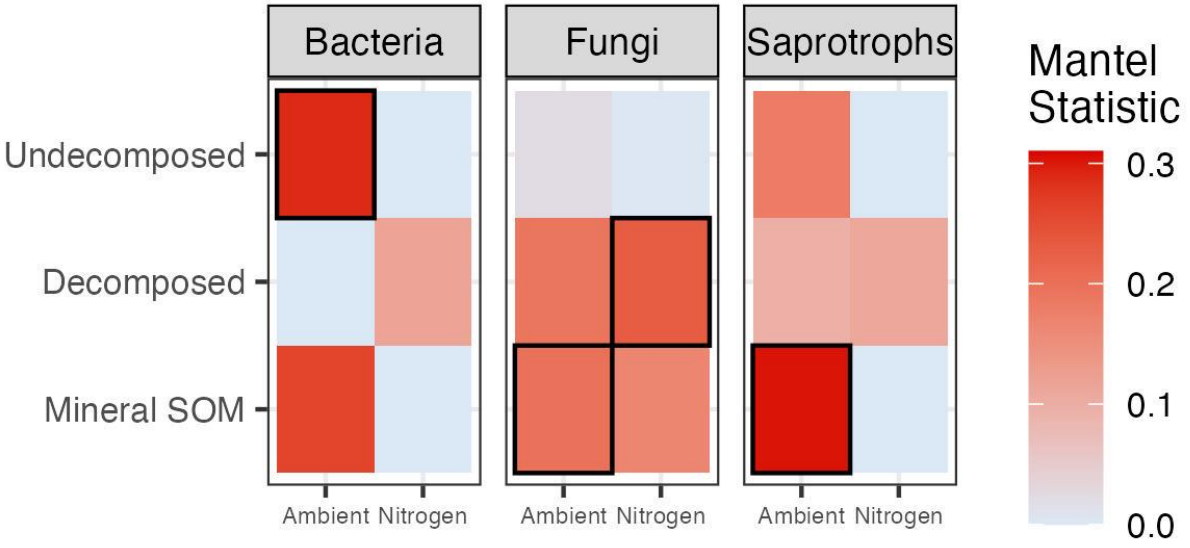


**Figure S4.** Mantel statistic (R) for the relationships between the bacterial (left), fungal (middle), and saprotrophic (right) communities and the biochemistry of the organic matter at each decomposition stage. Higher R values indicate stronger relationships (red). Significant relationships ($P \leq 0.1$) are outlined. See Table S13 for full Mantel statistics.

**Table S13.** Mantel test results comparing $log_2$-transformed Euclidean distance matrices of individual compound abundances and Hellinger-transformed Bray-Curtis distance matrices of microbial community OTU abundances. Significant P-values ($P \leq 0.1$) are presented in bold.

| | | | Mantel | P- | Upper quantiles of permutations (null model) | | | |
|---|---|---|---|---|---|---|---|---|
| Type | Treatment | Decomp Stage | Statistic (R) | value | 90% | 95% | 97.5% | 99% |
| Bacteria (16S) | Ambient | Undecomposed | **0.288700** | **0.069** | 0.255 | 0.314 | 0.360 | 0.406 |
| | | Decomposed | -0.001153 | 0.476 | 0.245 | 0.339 | 0.428 | 0.474 |
| | | Mineral SOM | 0.2607 | 0.123 | 0.296 | 0.375 | 0.426 | 0.467 |
| | Nitrogen | Undecomposed | -0.2065 | 0.912 | 0.227 | 0.278 | 0.328 | 0.371 |
| | | Decomposed | 0.117 | 0.211 | 0.278 | 0.376 | 0.441 | 0.518 |
| | | Mineral SOM | -0.1325 | 0.720 | 0.261 | 0.321 | 0.384 | 0.425 |
| Fungi (28S) | Ambient | Undecomposed | 0.02193 | 0.455 | 0.203 | 0.251 | 0.299 | 0.341 |
| | | Decomposed | 0.1921 | 0.107 | 0.200 | 0.237 | 0.273 | 0.326 |
| | | Mineral SOM | **0.1981** | **0.091** | 0.187 | 0.239 | 0.292 | 0.328 |
| | Nitrogen | Undecomposed | 0.002091 | 0.473 | 0.161 | 0.210 | 0.243 | 0.290 |
| | | Decomposed | **0.2292** | **0.041** | 0.175 | 0.208 | 0.246 | 0.298 |
| | | Mineral SOM | 0.1662 | 0.105 | 0.170 | 0.216 | 0.243 | 0.298 |
| Saprotrophs | Ambient | Undecomposed | 0.1829 | 0.148 | 0.230 | 0.275 | 0.316 | 0.359 |
| | | Decomposed | 0.09608 | 0.290 | 0.229 | 0.284 | 0.363 | 0.413 |
| | | Mineral SOM | **0.3031** | **0.064** | 0.240 | 0.319 | 0.368 | 0.411 |
| | Nitrogen | Undecomposed | -0.01454 | 0.518 | 0.229 | 0.281 | 0.335 | 0.382 |
| | | Decomposed | 0.1114 | 0.242 | 0.256 | 0.336 | 0.408 | 0.467 |
| | | Mineral SOM | -0.1032 | 0.681 | 0.281 | 0.347 | 0.382 | 0.449 |

**Table S14.** Mineral soil $log_2$-transformed individual compound abundance vectors fit to Hellinger-transformed bacterial (16S) community db-RDA ordination. Compounds with significant P-values ($P \leq 0.1$) are presented in bold.

| Compound | Source Class | $R^2$ | P-value |
|---|---|---|---|
| Phenol, 3,4-dimethyl- | Aromatic | 0.1118 | 0.283 |
| Benzene, 1,2-diethyl- | Aromatic | 0.0281 | 0.739 |
| Benzaldehyde | Aromatic | 0.0654 | 0.501 |
| Oxirane, ethenyl- | Aromatic | 0.0311 | 0.735 |
| Acetophenone | Aromatic | 0.0266 | 0.739 |
| Benzene, 1,2,3-trimethyl- | Aromatic | 0.0088 | 0.929 |
| Benzene, 1-ethenyl-3-methyl- | Aromatic | 0.0301 | 0.753 |
| Benzene, (1-methylethyl)- | Aromatic | 0.0585 | 0.527 |
| **Benzofuran** | **Aromatic** | **0.2971** | **0.021** |
| Benzene | Aromatic | 0.0427 | 0.634 |
| Fluorene | Aromatic | 0.1333 | 0.210 |
| Naphthalene | Aromatic | 0.0116 | 0.891 |
| Benzene, 2-propenyl- | Aromatic | 0.0673 | 0.462 |
| m-xylene | Aromatic | 0.0502 | 0.571 |
| Benzene, propyl- | Aromatic | 0.0024 | 0.970 |
| Benzene, hexyl- | Aromatic | 0.0410 | 0.632 |
| Benzene | Aromatic | 0.0091 | 0.904 |
| Benzene, butyl- | Aromatic | 0.0245 | 0.760 |
| **Phenol, 3-methyl-** | **Aromatic** | **0.2412** | **0.063** |
| Naphthalene, 1,2-dihydro-4-methyl- | Aromatic | 0.0260 | 0.758 |
| Benzene, (1,3-dimethylbutyl)- | Aromatic | 0.1180 | 0.263 |
| Biphenyl | Aromatic | 0.0089 | 0.914 |
| Anthracene | Aromatic | 0.0307 | 0.720 |
| Benzene, 1,2,3,4-tetramethyl- | Aromatic | 0.0924 | 0.372 |
| 2H-1-Benzopyran-2-one | Aromatic | 0.0557 | 0.636 |
| Phenol, 2-methoxy-4-(1Z)-1-propen-1-yl- (cis-Isoeugenol) | Lignin | 0.0324 | 0.704 |
| **Ethanone, 1-(4-hydroxy-3-methoxyphenyl)- (Acetovanillone)** | **Lignin** | **0.3173** | **0.029** |
| Phenol, 2-methoxy-4-(2-propenyl)- (4-Eugenol) | Lignin | 0.0554 | 0.570 |
| Benzaldehyde, 4-hydroxy-3-methoxy- (Vanillin) | Lignin | 0.0598 | 0.501 |
| **2-Propanone, 1-(4-hydroxy-3-methoxyphenyl)- (Guaiacylacetone)** | **Lignin** | **0.4068** | **0.017** |
| **Phenol, 2,6-dimethoxy-4-(2-propenyl)- (Methoxyeugenol)** | **Lignin** | **0.5126** | **0.004** |
| Benzeneacetic acid, 4-hydroxy-3-methoxy- (Homovanillic acid) | Lignin | 0.4366 | 0.020 |
| Phenol, 4-ethyl-2-methoxy- (Ethylguaiacol) | Lignin | 0.0275 | 0.758 |
| Phenol, 2,6-dimethoxy- (Syringol) | Lignin | 0.0102 | 0.904 |
| **Phenol, 2-methoxy-4-propyl- (4-Guaiacylpropane)** | **Lignin** | **0.4187** | **0.019** |
| **Ethanone, 1-(4-hydroxy-3,5-dimethoxyphenyl)- (Acetosyringone)** | **Lignin** | **0.2798** | **0.029** |
| Ethylphenol | Lignin | 0.0939 | 0.366 |
| **Phenol, 4-ethyl-2,6-dimethoxy- (Ethylsyringol)** | **Lignin** | **0.4365** | **0.045** |
| Phenol, 2-methoxy- (Guaiacol) | Lignin | 0.1124 | 0.204 |
| Benzoic acid, 4-hydroxy-3-methoxy-Vanillic acid) | Lignin | 0.0517 | 0.692 |
| **Ethanone, 1-(3,4-dimethoxyphenyl)-** | Lignin | **0.2806** | **0.036** |
| Benzoic acid, 4-hydroxy-3-methoxy-, methyl ester (Vanillic Acid, methyl est" | Lignin | 0.0869 | 0.365 |
| Benzene, 4-ethenyl-1,2-dimethoxy- | Lignin | 0.0940 | 0.380 |
| Benzene, 1-methoxy-4-methyl- | Lignin | 0.1171 | 0.229 |
| **3,4-Dimethoxytoluene** | **Lignin** | **0.3115** | **0.045** |
| **Benzene, 1-ethenyl-4-methoxy-** | **Lignin** | **0.3466** | **0.008** |

| | | | |
|---|---|---|---|
| **Hexadecanoic acid, methyl ester (Palmitic acid-C16)** | **Lipid** | **0.2577** | **0.055** |
| n-Docosane (C22) | Lipid | 0.0940 | 0.380 |
| 1-Butyne, 3,3-dimethyl- | Lipid | 0.0012 | 0.988 |
| 3-Decene | Lipid | 0.0296 | 0.726 |
| 7-Tetradecene | Lipid | 0.0002 | 0.998 |
| Propene | Lipid | 0.0839 | 0.375 |
| Isocrotonic acid | Lipid | 0.0589 | 0.492 |
| 2-Butenedioic acid, 2-methyl-, (E)- | Lipid | 0.0423 | 0.681 |
| 1-Hexene, 3-methyl- | Lipid | 0.0244 | 0.764 |
| 1-Heptene | Lipid | 0.0730 | 0.457 |
| n-Nonane | Lipid | 0.0300 | 0.745 |
| 1,3-Butadiene | Lipid | 0.0364 | 0.663 |
| C14_alkene_3 | Lipid | 0.0804 | 0.394 |
| 1-Pentene, 3-ethyl-2-methyl- | Lipid | 0.1815 | 0.118 |
| n-Undecane | Lipid | 0.0392 | 0.666 |
| n-Pentadecane | Lipid | 0.0261 | 0.765 |
| 1,3-Octadiene | Lipid | 0.0429 | 0.648 |
| n-Octane | Lipid | 0.0967 | 0.340 |
| n-Heptane | Lipid | 0.0748 | 0.437 |
| n-Tetracosane | Lipid | 0.0940 | 0.380 |
| 1-Hexadecene | Lipid | 0.1146 | 0.285 |
| n-Dodecane | Lipid | 0.0018 | 0.978 |
| Hex-2-yn-4-one, 2-methyl- | Lipid | 0.0008 | 0.990 |
| n-Nonacosane (C29) | Lipid | 0.1757 | 0.129 |
| **n-Heptacosane (C27)** | Lipid | **0.2022** | **0.100** |
| n-Tridecane | Lipid | 0.0296 | 0.725 |
| n-Heptadecane | Lipid | 0.0647 | 0.507 |
| **n-Decane** | Lipid | **0.2289** | **0.067** |
| n-Hexadecane | Lipid | 0.0067 | 0.959 |
| n-Tricosane (C23) | Lipid | 0.0628 | 0.492 |
| n-Tetradecane | Lipid | 0.1519 | 0.188 |
| n-Hexacosane (C26) | Lipid | 0.0201 | 0.817 |
| Hepta-2,4-dienoic acid, methyl ester | Lipid | 0.0101 | 0.888 |
| n-Heneicosane (C21) | Lipid | 0.0032 | 1.000 |
| **Pyrazolo[5,1-c][1,2,4]benzotriazin-8-ol** | **N-Bearing** | **0.5926** | **0.001** |
| N-Butyl-tert-butylamine | N-Bearing | 0.0331 | 0.720 |
| 1H-Tetrazole, 1-methyl- | N-Bearing | 0.0532 | 0.556 |
| **3-Pyridinol** | **N-Bearing** | **0.2320** | **0.067** |
| 1H-Pyrrole-2-carboxaldehyde | N-Bearing | 0.0412 | 0.635 |
| 2,5-Furandione, 3-methyl- | N-Bearing | 0.0528 | 0.552 |
| Aniline | N-Bearing | 0.0325 | 0.713 |
| 1H-Pyrrole, 3-methyl- | N-Bearing | 0.0429 | 0.628 |
| Pyridine 3-methyl | N-Bearing | 0.0983 | 0.319 |
| Pyridine, 4-methoxy- | N-Bearing | 0.0683 | 0.454 |
| p-Aminotoluene | N-Bearing | 0.0879 | 0.376 |
| Hexanedinitrile | N-Bearing | 0.0341 | 0.719 |
| 1H-Pyrrole-2-carboxaldehyde, 1-methyl- | N-Bearing | 0.0442 | 0.620 |
| Pyridine, 2-ethyl | N-Bearing | 0.0846 | 0.398 |
| 3-Methylpyridazine | N-Bearing | 0.1894 | 0.124 |
| 3-Phenylpyridine | N-Bearing | 0.1400 | 0.186 |
| Propane, 2-nitro- | N-Bearing | 0.0136 | 0.876 |
| 4-Pyridinecarboxaldehyde | N-Bearing | 0.0339 | 0.703 |
| 2-Pyridinealdehyde | N-Bearing | 0.0567 | 0.531 |
| 4(1H)-Pyridinone, 2,3-dihydro-1-methyl- | N-Bearing | 0.0055 | 0.943 |
| Alpha-amino-gamma-butyrolactone | N-Bearing | 0.1267 | 0.187 |
| Pyrimidine | N-Bearing | 0.1382 | 0.183 |

| | | | |
|---|---|---|---|
| Piperidine-2,5-dione | N-Bearing | 0.0734 | 0.487 |
| Acetamide, N-(2,4-dihydroxyphenyl)- | N-Bearing | 0.0650 | 0.502 |
| **4-Pyridinecarbonitrile** | **N-Bearing** | **0.2651** | **0.046** |
| Diethyltoluamide (DEET) | N-Bearing | 0.0593 | 0.518 |
| 5H-1-Pyrindine | N-Bearing | 0.0510 | 0.573 |
| 1,4-Benzenediamine | N-Bearing | 0.0326 | 0.671 |
| 5-Dimethylaminopyrimidine | N-Bearing | 0.0180 | 0.823 |
| Ethanone, 1-(1-methyl-1H-pyrrol-2-yl)- | N-Bearing | 0.0164 | 0.854 |
| Acetamide, N-hydroxy | N-Bearing | 0.0275 | 0.758 |
| 2-(N-Methyl-N-ethylamino)phenol | N-Bearing | 0.0461 | 0.617 |
| Cyanamide, dimethyl- | N-Bearing | 0.0032 | 1.000 |
| 2-Pyrimidinamine | N-Bearing | 0.0032 | 1.000 |
| Methylaminoacetonitrile | N-Bearing | 0.0593 | 0.554 |
| 2-Amino-4-methylpyrimidine | N-Bearing | 0.1643 | 0.157 |
| 3-Pyridinecarbonitrile | N-Bearing | 0.0409 | 0.630 |
| 4-Amino-2(1H)-pyridinone | N-Bearing | 0.0275 | 0.758 |
| N-Isopropyl-4-piperidone | N-Bearing | 0.0275 | 0.758 |
| Nicotinyl Alcohol | N-Bearing | 0.0275 | 0.758 |
| Phenol | Phenol | 0.1915 | 0.129 |
| Phenol, 4-methyl- | Phenol | 0.1842 | 0.129 |
| **Benzene, 1,4-dimethoxy-** | **Phenol** | **0.4442** | **0.016** |
| Benzene, 1-ethyl-4-methoxy- | Phenol | 0.1086 | 0.318 |
| 3-Furaldehyde | Polysaccharide | 0.0238 | 0.786 |
| Levoglucosan | Polysaccharide | 0.0366 | 0.702 |
| 2(3H)-Furanone, 5-methyl- | Polysaccharide | 0.0373 | 0.673 |
| **2-Cyclopenten-1-one, 2-hydroxy-3-methyl-** | **Polysaccharide** | **0.2540** | **0.046** |
| Furfural, 5-methyl- | Polysaccharide | 0.0353 | 0.693 |
| Benzofuran, 2,3-dihydro- | Polysaccharide | 0.0561 | 0.585 |
| Acetic anhydride | Polysaccharide | 0.0646 | 0.489 |
| 4H-Pyran-4-one, 3-hydroxy-2-methyl- | Polysaccharide | 0.0246 | 0.752 |
| 2-Furanmethanol | Polysaccharide | 0.1267 | 0.187 |
| Furan, 2,4-dimethyl- | Polysaccharide | 0.0027 | 0.974 |
| 2-Cyclopenten-1-one, 2-methyl- | Polysaccharide | 0.0231 | 0.777 |
| 2-Acetylfuran | Polysaccharide | 0.0488 | 0.599 |
| 2H-Pyran-2-one | Polysaccharide | 0.0578 | 0.508 |
| 1,2-Cyclohexanedione | Polysaccharide | 0.0651 | 0.518 |
| Butanal, 2-methyl- | Polysaccharide | 0.0623 | 0.489 |
| 2-Cyclopenten-1-one, 2,3-dimethyl- | Polysaccharide | 0.0341 | 0.688 |
| Furan, 2,5-dimethyl- | Polysaccharide | 0.0440 | 0.604 |
| Levoglucosenone | Polysaccharide | 0.0514 | 0.565 |
| 2(5H)-Furanone, 5-methyl- | Polysaccharide | 0.0789 | 0.400 |
| Furan, 2-ethyl-5-methyl- | Polysaccharide | 0.0277 | 0.725 |
| Benzofuran, 2-methyl- | Polysaccharide | 0.0636 | 0.494 |
| Furan, 2,3,5-trimethyl- | Polysaccharide | 0.0251 | 0.724 |
| Vinylfuran | Polysaccharide | 0.0628 | 0.485 |
| Cyclopent-2-ene-1-one, 2,3,4-trimethyl- | Polysaccharide | 0.0579 | 0.560 |
| 1,2-Cyclopentanedione, 3-methyl- | Polysaccharide | 0.0275 | 0.758 |
| PENTANAL | Polysaccharide | 0.0747 | 0.452 |
| Furan, 2-ethyl- | Polysaccharide | 0.0637 | 0.504 |
| 1,2-Cyclopentanedione | Polysaccharide | 0.0345 | 0.689 |
| Cyclopropanecarboxaldehyde, methylene- | Polysaccharide | 0.0087 | 0.910 |
| Cyclopentanone | Polysaccharide | 0.0611 | 0.512 |
| 3-Acetamidofuran | Polysaccharide | 0.0724 | 0.432 |
| Furfural | Polysaccharide | 0.0988 | 0.318 |
| 5-Ethyl-2-furaldehyde | Polysaccharide | 0.0101 | 0.883 |
| 2(3H)-Benzofuranone, 3-methyl- | Polysaccharide | 0.0423 | 0.681 |

| | | | |
|---|---|---|---|
| Indole | Protein | 0.1154 | 0.292 |
| Styrene | Protein | 0.0360 | 0.689 |
| Benzyl nitrile | Protein | 0.0091 | 0.909 |
| 3-Methylindole | Protein | 0.0161 | 0.854 |
| Ethylbenzene | Protein | 0.0393 | 0.649 |
| Pyridine | Protein | 0.0479 | 0.584 |
| 4-Pyridinamine | Protein | 0.1577 | 0.193 |
| 1H-Pyrrole, 1-methyl- | Protein | 0.0408 | 0.670 |
| Pyridine, 3,5-dimethyl- | Protein | 0.0138 | 0.852 |
| Pyrrole | Protein | 0.0403 | 0.654 |
| Benzonitrile | Protein | 0.0340 | 0.710 |
| **1 H-Pyrrole, 2-ethyl-** | **Protein** | **0.3200** | **0.013** |
| Pyridine | Protein | 0.0185 | 0.832 |
| Benzenepropanenitrile | Protein | 0.1056 | 0.265 |
| Mequinol | Unspecified | 0.0833 | 0.416 |
| Spiro[2.4]hepta-4,6-diene | Unspecified | 0.0507 | 0.568 |
| **(ISTD) 3-Ethoxy-4-methoxybenzaldehyde** | **Unspecified** | **0.4365** | **0.045** |
| Hydroquinone | Unspecified | 0.0272 | 0.777 |
| 1,3,5-Cycloheptatriene | Unspecified | 0.0669 | 0.478 |
| Methanesulfonic acid, methyl ester | Unspecified | 0.0267 | 0.740 |
| Benzene, 1,4-dimethoxy-2,3,5,6-tetramethyl- | Unspecified | 0.1267 | 0.187 |
| Pyruvaldehyde | Unspecified | 0.1268 | 0.249 |
| Monobenzone | Unspecified | 0.0096 | 0.911 |
| C8_H16 | Unspecified | 0.0334 | 0.701 |
| 1-Undecanol | Unspecified | 0.0041 | 0.966 |
| Indane | Unspecified | 0.0458 | 0.630 |
| Dimethylbenzofuran | Unspecified | 0.0177 | 0.836 |
| 1H-Inden-1-one, 2,3-dihydro- | Unspecified | 0.0220 | 0.797 |
| Phenanthrene | Unspecified | 0.0015 | 0.989 |
| 2,3,6-Trimethylnaphthalene | Unspecified | 0.0587 | 0.494 |
| **Trimethylphenol** | **Unspecified** | **0.3254** | **0.025** |
| Vinyl crotonate | Unspecified | 0.0275 | 0.758 |
| C9_H8 | Unspecified | 0.0009 | 0.989 |
| Toluene | Unspecified | 0.0052 | 0.952 |
| 3-Penten-2-one, (E)- | Unspecified | 0.0551 | 0.547 |
| 4-Hydroxy-2-methylacetophenone | Unspecified | 0.0041 | 0.951 |
| **Benzoic acid, 4-hydroxy-3,5-dimethoxy-, hydrazide** | **Unspecified** | **0.4207** | **0.005** |
| 1-Propene, 3-azido- | Unspecified | 0.0464 | 0.626 |
| **2-Pentanone** | **Unspecified** | **0.4365** | **0.045** |
| Resorcinol (Dihydroxybenzene) | Unspecified | 0.0324 | 0.706 |
| Acenaphthene | Unspecified | 0.0193 | 0.805 |
| Ethanone, 1-(3-hydroxy-4-methoxyphenyl)- | Unspecified | 0.0738 | 0.476 |
| 2-Propenoic acid, ethenyl ester | Unspecified | 0.0042 | 0.959 |
| **2-Methoxy-5-methylphenol** | **Unspecified** | **0.4365** | **0.045** |
| 2-Propen-1-ol | Unspecified | 0.0517 | 0.692 |
| C11_H12 | Unspecified | 0.1372 | 0.197 |
| (E)-1,3-Butadien-1-ol | Unspecified | 0.0421 | 0.667 |
| Thiophene | Unspecified | 0.1267 | 0.187 |
| Benzene, heptyl- | Unspecified | 0.0394 | 0.677 |
| 3-Methylthiophene-2-carbonitrile | Unspecified | 0.1160 | 0.226 |
| Fluoranthene | Unspecified | 0.0367 | 0.710 |
| Pyrene | Unspecified | 0.0055 | 0.932 |
| 7-Methylindan-1-one | Unspecified | 0.0269 | 0.764 |
| Acenaphthylene | Unspecified | 0.0517 | 0.692 |
| **Cyclobut-1-enylmethanol** | **Unspecified** | **0.4365** | **0.045** |
| **2-Methylthiophene** | **Unspecified** | **0.4365** | **0.045** |

| | | | |
|---|---|---|---|
| Isobutyl nitrite | Unspecified | 0.1267 | 0.187 |
| Phenol, 2-methoxy-5-(1-propenyl)-, (E)- | Unspecified | 0.0423 | 0.681 |

**Table S15.** Mineral soil $log_2$-transformed individual compound abundance vectors fit to Hellinger-transformed fungal (28S) community db-RDA ordination. Compounds with significant P-values ($P \leq 0.1$) are presented in bold.

| Compound | Source Class | $R^2$ | P-value |
|---|---|---|---|
| Phenol, 3,4-dimethyl- | Aromatic | 0.1018 | 0.347 |
| Benzene, 1,2-diethyl- | Aromatic | 0.0665 | 0.454 |
| Benzaldehyde | Aromatic | 0.0023 | 0.984 |
| Oxirane, ethenyl- | Aromatic | 0.0771 | 0.455 |
| Acetophenone | Aromatic | 0.0565 | 0.539 |
| Benzene, 1,2,3-trimethyl- | Aromatic | 0.1046 | 0.318 |
| **Benzene, 1-ethenyl-3-methyl-** | **Aromatic** | **0.3458** | **0.012** |
| Benzene, (1-methylethyl)- | Aromatic | 0.0797 | 0.433 |
| **Benzofuran** | **Aromatic** | **0.2817** | **0.025** |
| Benzene | Aromatic | 0.0332 | 0.713 |
| Fluorene | Aromatic | 0.0767 | 0.430 |
| Naphthalene | Aromatic | 0.1888 | 0.109 |
| Benzene, 2-propenyl- | Aromatic | 0.1668 | 0.162 |
| m-xylene | Aromatic | 0.0904 | 0.399 |
| Benzene, propyl- | Aromatic | 0.1999 | 0.102 |
| **Benzene, hexyl-** | **Aromatic** | **0.2145** | **0.085** |
| Benzene | Aromatic | 0.0895 | 0.391 |
| **Benzene, butyl-** | **Aromatic** | **0.2029** | **0.098** |
| **Phenol, 3-methyl-** | **Aromatic** | **0.2135** | **0.075** |
| Naphthalene, 1,2-dihydro-4-methyl- | Aromatic | 0.1452 | 0.194 |
| Benzene, (1,3-dimethylbutyl)- | Aromatic | 0.1362 | 0.232 |
| Biphenyl | Aromatic | 0.0893 | 0.396 |
| Anthracene | Aromatic | 0.0003 | 0.997 |
| **Benzene, 1,2,3,4-tetramethyl-** | **Aromatic** | **0.2942** | **0.021** |
| 2H-1-Benzopyran-2-one | Aromatic | 0.1285 | 0.292 |
| **Phenol, 2-methoxy-4-(1Z)-1-propen-1-yl- (cis-Isoeugenol)** | **Lignin** | **0.2204** | **0.085** |
| **Ethanone, 1-(4-hydroxy-3-methoxyphenyl)- (Acetovanillone)** | **Lignin** | **0.2698** | **0.036** |
| Phenol, 2-methoxy-4-(2-propenyl)- (4-Eugenol) | Lignin | 0.1928 | 0.123 |
| Benzaldehyde, 4-hydroxy-3-methoxy- (Vanillin) | Lignin | 0.0443 | 0.634 |
| **2-Propanone, 1-(4-hydroxy-3-methoxyphenyl)- (Guaiacylacetone)** | **Lignin** | **0.3564** | **0.006** |
| **Phenol, 2,6-dimethoxy-4-(2-propenyl)- (Methoxyeugenol)** | **Lignin** | **0.6315** | **0.001** |
| **Benzeneacetic acid, 4-hydroxy-3-methoxy- (Homovanillic acid)** | **Lignin** | **0.3192** | **0.007** |
| Phenol, 4-ethyl-2-methoxy- (Ethylguaiacol) | Lignin | 0.0075 | 0.874 |
| Phenol, 2,6-dimethoxy- (Syringol) | Lignin | 0.1985 | 0.104 |
| **Phenol, 2-methoxy-4-propyl- (4-Guaiacylpropane)** | **Lignin** | **0.2414** | **0.038** |
| **Ethanone, 1-(4-hydroxy-3,5-dimethoxyphenyl)- (Acetosyringone)** | **Lignin** | **0.4731** | **0.004** |
| Ethylphenol | Lignin | 0.0309 | 0.709 |
| **Phenol, 4-ethyl-2,6-dimethoxy- (Ethylsyringol)** | **Lignin** | **0.2902** | **0.045** |
| Phenol, 2-methoxy- (Guaiacol) | Lignin | 0.0252 | 0.788 |
| Benzoic acid, 4-hydroxy-3-methoxy-Vanillic acid) | Lignin | 0.0011 | 1.000 |
| **Ethanone, 1-(3,4-dimethoxyphenyl)-** | **Lignin** | **0.5407** | **0.002** |
| Benzoic acid, 4-hydroxy-3-methoxy-, methyl ester (Vanillic Acid, methyl est" | Lignin | 0.0778 | 0.438 |
| **Benzene, 4-ethenyl-1,2-dimethoxy-** | **Lignin** | **0.2441** | **0.077** |
| Benzene, 1-methoxy-4-methyl- | Lignin | 0.1807 | 0.109 |
| 3,4-Dimethoxytoluene | Lignin | 0.1896 | 0.114 |
| Benzene, 1-ethenyl-4-methoxy- | Lignin | 0.0383 | 0.687 |

| | | | |
|---|---|---|---|
| Hexadecanoic acid, methyl ester (Palmitic acid-C16) | Lipid | 0.0880 | 0.411 |
| **n-Docosane (C22)** | **Lipid** | **0.2441** | **0.077** |
| 1-Butyne, 3,3-dimethyl- | Lipid | 0.0353 | 0.672 |
| 3-Decene | Lipid | 0.2057 | 0.111 |
| 7-Tetradecene | Lipid | 0.1253 | 0.271 |
| Propene | Lipid | 0.0165 | 0.866 |
| Isocrotonic acid | Lipid | 0.0471 | 0.631 |
| 2-Butenedioic acid, 2-methyl-, (E)- | Lipid | 0.0752 | 0.485 |
| 1-Hexene, 3-methyl- | Lipid | 0.0429 | 0.638 |
| 1-Heptene | Lipid | 0.0268 | 0.748 |
| **n-Nonane** | **Lipid** | **0.3118** | **0.021** |
| 1,3-Butadiene | Lipid | 0.0690 | 0.456 |
| **C14_alkene_3** | **Lipid** | **0.2313** | **0.054** |
| 1-Pentene, 3-ethyl-2-methyl- | Lipid | 0.0455 | 0.624 |
| **n-Undecane** | **Lipid** | **0.2176** | **0.098** |
| n-Pentadecane | Lipid | 0.1238 | 0.274 |
| 1,3-Octadiene | Lipid | 0.0106 | 0.891 |
| **n-Octane** | **Lipid** | **0.2201** | **0.086** |
| n-Heptane | Lipid | 0.1428 | 0.175 |
| **n-Tetracosane** | **Lipid** | **0.2441** | **0.077** |
| 1-Hexadecene | Lipid | 0.1682 | 0.154 |
| n-Dodecane | Lipid | 0.0667 | 0.476 |
| Hex-2-yn-4-one, 2-methyl- | Lipid | 0.0256 | 0.797 |
| n-Nonacosane (C29) | Lipid | 0.0347 | 0.700 |
| **n-Heptacosane (C27)** | **Lipid** | **0.2215** | **0.078** |
| n-Tridecane | Lipid | 0.1369 | 0.197 |
| n-Heptadecane | Lipid | 0.0863 | 0.388 |
| **n-Decane** | **Lipid** | **0.3227** | **0.021** |
| n-Hexadecane | Lipid | 0.0577 | 0.674 |
| n-Tricosane (C23) | Lipid | 0.0900 | 0.387 |
| n-Tetradecane | Lipid | 0.0550 | 0.551 |
| n-Hexacosane (C26) | Lipid | 0.0536 | 0.577 |
| Hepta-2,4-dienoic acid, methyl ester | Lipid | 0.0162 | 0.853 |
| n-Heneicosane (C21) | Lipid | 0.0026 | 0.946 |
| **Pyrazolo[5,1-c][1,2,4]benzotriazin-8-ol** | **N-Bearing** | **0.4244** | **0.007** |
| N-Butyl-tert-butylamine | N-Bearing | 0.0480 | 0.581 |
| 1H-Tetrazole, 1-methyl- | N-Bearing | 0.0664 | 0.483 |
| **3-Pyridinol** | **N-Bearing** | **0.4035** | **0.001** |
| 1H-Pyrrole-2-carboxaldehyde | N-Bearing | 0.0263 | 0.785 |
| 2,5-Furandione, 3-methyl- | N-Bearing | 0.0084 | 0.927 |
| Aniline | N-Bearing | 0.0878 | 0.386 |
| 1H-Pyrrole, 3-methyl- | N-Bearing | 0.1564 | 0.165 |
| Pyridine 3-methyl | N-Bearing | 0.0169 | 0.829 |
| Pyridine, 4-methoxy- | N-Bearing | 0.0238 | 0.788 |
| **p-Aminotoluene** | **N-Bearing** | **0.2018** | **0.095** |
| Hexanedinitrile | N-Bearing | 0.0603 | 0.526 |
| 1H-Pyrrole-2-carboxaldehyde, 1-methyl- | N-Bearing | 0.0053 | 0.937 |
| Pyridine, 2-ethyl | N-Bearing | 0.0123 | 0.872 |
| 3-Methylpyridazine | N-Bearing | 0.0607 | 0.517 |
| **3-Phenylpyridine** | **N-Bearing** | **0.2086** | **0.081** |
| Propane, 2-nitro- | N-Bearing | 0.0982 | 0.333 |
| 4-Pyridinecarboxaldehyde | N-Bearing | 0.0600 | 0.536 |
| 2-Pyridinealdehyde | N-Bearing | 0.0160 | 0.857 |
| 4(1H)-Pyridinone, 2,3-dihydro-1-methyl- | N-Bearing | 0.0529 | 0.576 |
| Alpha-amino-gamma-butyrolactone | N-Bearing | 0.0058 | 0.919 |
| Pyrimidine | N-Bearing | 0.0011 | 0.993 |

| | | | |
|---|---|---|---|
| Piperidine-2,5-dione | N-Bearing | 0.0436 | 0.624 |
| Acetamide, N-(2,4-dihydroxyphenyl)- | N-Bearing | 0.1287 | 0.244 |
| 4-Pyridinecarbonitrile | N-Bearing | 0.1629 | 0.159 |
| **Diethyltoluamide (DEET)** | **N-Bearing** | **0.3100** | **0.015** |
| 5H-1-Pyrindine | N-Bearing | 0.1344 | 0.218 |
| 1,4-Benzenediamine | N-Bearing | 0.0548 | 0.559 |
| 5-Dimethylaminopyrimidine | N-Bearing | 0.1259 | 0.255 |
| Ethanone, 1-(1-methyl-1H-pyrrol-2-yl)- | N-Bearing | 0.0027 | 0.970 |
| Acetamide, N-hydroxy | N-Bearing | 0.0075 | 0.874 |
| 2-(N-Methyl-N-ethylamino)phenol | N-Bearing | 0.0348 | 0.691 |
| Cyanamide, dimethyl- | N-Bearing | 0.0026 | 0.946 |
| 2-Pyrimidinamine | N-Bearing | 0.0026 | 0.946 |
| Methylaminoacetonitrile | N-Bearing | 0.0404 | 0.758 |
| 2-Amino-4-methylpyrimidine | N-Bearing | 0.0491 | 0.608 |
| 3-Pyridinecarbonitrile | N-Bearing | 0.0741 | 0.441 |
| 4-Amino-2(1H)-pyridinone | N-Bearing | 0.0075 | 0.874 |
| N-Isopropyl-4-piperidone | N-Bearing | 0.0075 | 0.874 |
| Nicotinyl Alcohol | N-Bearing | 0.0075 | 0.874 |
| Phenol | Phenol | 0.1186 | 0.281 |
| Phenol, 4-methyl- | Phenol | 0.0925 | 0.341 |
| **Benzene, 1,4-dimethoxy-** | **Phenol** | **0.3266** | **0.008** |
| Benzene, 1-ethyl-4-methoxy- | Phenol | 0.1107 | 0.309 |
| 3-Furaldehyde | Polysaccharide | 0.0397 | 0.638 |
| Levoglucosan | Polysaccharide | 0.0989 | 0.337 |
| 2(3H)-Furanone, 5-methyl- | Polysaccharide | 0.0756 | 0.429 |
| 2-Cyclopenten-1-one, 2-hydroxy-3-methyl- | Polysaccharide | 0.0600 | 0.515 |
| Furfural, 5-methyl- | Polysaccharide | 0.0482 | 0.583 |
| Benzofuran, 2,3-dihydro- | Polysaccharide | 0.1454 | 0.211 |
| Acetic anhydride | Polysaccharide | 0.0094 | 0.909 |
| 4H-Pyran-4-one, 3-hydroxy-2-methyl- | Polysaccharide | 0.1271 | 0.259 |
| 2-Furanmethanol | Polysaccharide | 0.0058 | 0.919 |
| Furan, 2,4-dimethyl- | Polysaccharide | 0.0408 | 0.619 |
| 2-Cyclopenten-1-one, 2-methyl- | Polysaccharide | 0.1666 | 0.144 |
| 2-Acetylfuran | Polysaccharide | 0.0624 | 0.489 |
| 2H-Pyran-2-one | Polysaccharide | 0.0448 | 0.622 |
| 1,2-Cyclohexanedione | Polysaccharide | 0.0125 | 0.891 |
| Butanal, 2-methyl- | Polysaccharide | 0.0557 | 0.568 |
| 2-Cyclopenten-1-one, 2,3-dimethyl- | Polysaccharide | 0.0124 | 0.881 |
| Furan, 2,5-dimethyl- | Polysaccharide | 0.0557 | 0.565 |
| Levoglucosenone | Polysaccharide | 0.0215 | 0.812 |
| 2(5H)-Furanone, 5-methyl- | Polysaccharide | 0.0266 | 0.779 |
| Furan, 2-ethyl-5-methyl- | Polysaccharide | 0.0184 | 0.820 |
| Benzofuran, 2-methyl- | Polysaccharide | 0.0131 | 0.886 |
| Furan, 2,3,5-trimethyl- | Polysaccharide | 0.0207 | 0.796 |
| Vinylfuran | Polysaccharide | 0.1200 | 0.250 |
| Cyclopent-2-ene-1-one, 2,3,4-trimethyl- | Polysaccharide | 0.0185 | 0.813 |
| 1,2-Cyclopentanedione, 3-methyl- | Polysaccharide | 0.0075 | 0.874 |
| PENTANAL | Polysaccharide | 0.1360 | 0.212 |
| Furan, 2-ethyl- | Polysaccharide | 0.0743 | 0.455 |
| 1,2-Cyclopentanedione | Polysaccharide | 0.0742 | 0.430 |
| Cyclopropanecarboxaldehyde, methylene- | Polysaccharide | 0.0498 | 0.673 |
| Cyclopentanone | Polysaccharide | 0.0272 | 0.774 |
| 3-Acetamidofuran | Polysaccharide | 0.1268 | 0.271 |
| Furfural | Polysaccharide | 0.1279 | 0.223 |
| 5-Ethyl-2-furaldehyde | Polysaccharide | 0.0860 | 0.377 |
| 2(3H)-Benzofuranone, 3-methyl- | Polysaccharide | 0.0752 | 0.485 |

| | | | |
|---|---|---|---|
| Indole | Protein | 0.1427 | 0.193 |
| Styrene | Protein | 0.0288 | 0.731 |
| Benzyl nitrile | Protein | 0.1532 | 0.169 |
| **3-Methylindole** | **Protein** | **0.1942** | **0.096** |
| Ethylbenzene | Protein | 0.0953 | 0.347 |
| Pyridine | Protein | 0.0420 | 0.633 |
| 4-Pyridinamine | Protein | 0.1791 | 0.124 |
| **1H-Pyrrole, 1-methyl-** | **Protein** | **0.2232** | **0.065** |
| Pyridine, 3,5-dimethyl- | Protein | 0.1102 | 0.264 |
| **Pyrrole** | **Protein** | **0.2342** | **0.063** |
| Benzonitrile | Protein | 0.0556 | 0.596 |
| 1 H-Pyrrole, 2-ethyl- | Protein | 0.0889 | 0.390 |
| Pyridine | Protein | 0.1338 | 0.222 |
| Benzenepropanenitrile | Protein | 0.0285 | 0.766 |
| Mequinol | Unspecified | 0.1416 | 0.193 |
| Spiro[2.4]hepta-4,6-diene | Unspecified | 0.1222 | 0.261 |
| **(ISTD) 3-Ethoxy-4-methoxybenzaldehyde** | **Unspecified** | **0.2902** | **0.045** |
| Hydroquinone | Unspecified | 0.0576 | 0.523 |
| 1,3,5-Cycloheptatriene | Unspecified | 0.0589 | 0.559 |
| Methanesulfonic acid, methyl ester | Unspecified | 0.0826 | 0.393 |
| Benzene, 1,4-dimethoxy-2,3,5,6-tetramethyl- | Unspecified | 0.0058 | 0.919 |
| Pyruvaldehyde | Unspecified | 0.0127 | 0.876 |
| Monobenzone | Unspecified | 0.0812 | 0.405 |
| C8_H16 | Unspecified | 0.1933 | 0.124 |
| **1-Undecanol** | **Unspecified** | **0.2184** | **0.088** |
| **Indane** | **Unspecified** | **0.3387** | **0.015** |
| Dimethylbenzofuran | Unspecified | 0.0111 | 0.890 |
| 1H-Inden-1-one, 2,3-dihydro- | Unspecified | 0.2029 | 0.104 |
| Phenanthrene | Unspecified | 0.0566 | 0.552 |
| 2,3,6-Trimethylnaphthalene | Unspecified | 0.1213 | 0.238 |
| Trimethylphenol | Unspecified | 0.1867 | 0.116 |
| Vinyl crotonate | Unspecified | 0.0075 | 0.874 |
| **C9_H8** | **Unspecified** | **0.2201** | **0.067** |
| Toluene | Unspecified | 0.0308 | 0.731 |
| 3-Penten-2-one, (E)- | Unspecified | 0.0248 | 0.749 |
| 4-Hydroxy-2-methylacetophenone | Unspecified | 0.0165 | 0.863 |
| **Benzoic acid, 4-hydroxy-3,5-dimethoxy-, hydrazide** | **Unspecified** | **0.5362** | **0.001** |
| 1-Propene, 3-azido- | Unspecified | 0.0786 | 0.423 |
| **2-Pentanone** | **Unspecified** | **0.2902** | **0.045** |
| Resorcinol (Dihydroxybenzene) | Unspecified | 0.1356 | 0.217 |
| Acenaphthene | Unspecified | 0.1140 | 0.295 |
| Ethanone, 1-(3-hydroxy-4-methoxyphenyl)- | Unspecified | 0.0763 | 0.441 |
| 2-Propenoic acid, ethenyl ester | Unspecified | 0.0221 | 0.795 |
| **2-Methoxy-5-methylphenol** | **Unspecified** | **0.2902** | **0.045** |
| 2-Propen-1-ol | Unspecified | 0.0011 | 1.000 |
| C11_H12 | Unspecified | 0.1494 | 0.206 |
| (E)-1,3-Butadien-1-ol | Unspecified | 0.0012 | 0.970 |
| Thiophene | Unspecified | 0.0058 | 0.919 |
| Benzene, heptyl- | Unspecified | 0.1220 | 0.277 |
| 3-Methylthiophene-2-carbonitrile | Unspecified | 0.0666 | 0.524 |
| Fluoranthene | Unspecified | 0.1236 | 0.277 |
| Pyrene | Unspecified | 0.0097 | 0.914 |
| 7-Methylindan-1-one | Unspecified | 0.0989 | 0.367 |
| Acenaphthylene | Unspecified | 0.0011 | 1.000 |
| **Cyclobut-1-enylmethanol** | **Unspecified** | **0.2902** | **0.045** |
| **2-Methylthiophene** | **Unspecified** | **0.2902** | **0.045** |

| | | | |
|---|---|---|---|
| Isobutyl nitrite | Unspecified | 0.0058 | 0.919 |
| Phenol, 2-methoxy-5-(1-propenyl)-, (E)- | Unspecified | 0.0752 | 0.485 |

**Table S16.** Mineral soil $log_2$-transformed individual compound abundance vectors fit to Hellinger-transformed saprotrophic community db-RDA ordination. Compounds with significant P-values ($P \leq 0.1$) are presented in bold.

| Compound | Source Class | $R^2$ | P-value |
|---|---|---|---|
| **Phenol, 3,4-dimethyl-** | **Aromatic** | **0.2147** | **0.067** |
| Benzene, 1,2-diethyl- | Aromatic | 0.0616 | 0.500 |
| Benzaldehyde | Aromatic | 0.1128 | 0.271 |
| Oxirane, ethenyl- | Aromatic | 0.0258 | 0.774 |
| Acetophenone | Aromatic | 0.0726 | 0.450 |
| Benzene, 1,2,3-trimethyl- | Aromatic | 0.0004 | 0.996 |
| Benzene, 1-ethenyl-3-methyl- | Aromatic | 0.0667 | 0.490 |
| Benzene, (1-methylethyl)- | Aromatic | 0.0890 | 0.345 |
| Benzofuran | Aromatic | 0.1264 | 0.256 |
| Benzene | Aromatic | 0.0464 | 0.614 |
| Fluorene | Aromatic | 0.0461 | 0.607 |
| Naphthalene | Aromatic | 0.0326 | 0.703 |
| Benzene, 2-propenyl- | Aromatic | 0.0534 | 0.557 |
| m-xylene | Aromatic | 0.0497 | 0.595 |
| Benzene, propyl- | Aromatic | 0.0099 | 0.896 |
| Benzene, hexyl- | Aromatic | 0.0601 | 0.491 |
| Benzene | Aromatic | 0.0035 | 0.958 |
| Benzene, butyl- | Aromatic | 0.0368 | 0.667 |
| **Phenol, 3-methyl-** | **Aromatic** | **0.3508** | **0.010** |
| Naphthalene, 1,2-dihydro-4-methyl- | Aromatic | 0.0091 | 0.914 |
| Benzene, (1,3-dimethylbutyl)- | Aromatic | 0.1417 | 0.200 |
| Biphenyl | Aromatic | 0.0326 | 0.702 |
| Anthracene | Aromatic | 0.0123 | 0.889 |
| Benzene, 1,2,3,4-tetramethyl- | Aromatic | 0.0392 | 0.667 |
| 2H-1-Benzopyran-2-one | Aromatic | 0.0941 | 0.447 |
| Phenol, 2-methoxy-4-(1Z)-1-propen-1-yl- (cis-Isoeugenol) | Lignin | 0.0641 | 0.521 |
| Ethanone, 1-(4-hydroxy-3-methoxyphenyl)- (Acetovanillone) | Lignin | 0.1897 | 0.110 |
| Phenol, 2-methoxy-4-(2-propenyl)- (4-Eugenol) | Lignin | 0.0720 | 0.447 |
| Benzaldehyde, 4-hydroxy-3-methoxy- (Vanillin) | Lignin | 0.0401 | 0.632 |
| **2-Propanone, 1-(4-hydroxy-3-methoxyphenyl)- (Guaiacylacetone)** | **Lignin** | **0.2850** | **0.014** |
| **Phenol, 2,6-dimethoxy-4-(2-propenyl)- (Methoxyeugenol)** | **Lignin** | **0.3602** | **0.005** |
| **Benzeneacetic acid, 4-hydroxy-3-methoxy- (Homovanillic acid)** | **Lignin** | **0.2293** | **0.014** |
| Phenol, 4-ethyl-2-methoxy- (Ethylguaiacol) | Lignin | 0.0332 | 0.762 |
| Phenol, 2,6-dimethoxy- (Syringol) | Lignin | 0.0011 | 0.988 |
| **Phenol, 2-methoxy-4-propyl- (4-Guaiacylpropane)** | **Lignin** | **0.2056** | **0.085** |
| **Ethanone, 1-(4-hydroxy-3,5-dimethoxyphenyl)- (Acetosyringone)** | **Lignin** | **0.3755** | **0.007** |
| Ethylphenol | Lignin | 0.1377 | 0.216 |
| **Phenol, 4-ethyl-2,6-dimethoxy- (Ethylsyringol)** | **Lignin** | **0.2108** | **0.045** |
| Phenol, 2-methoxy- (Guaiacol) | Lignin | 0.0146 | 0.875 |
| Benzoic acid, 4-hydroxy-3-methoxy-Vanillic acid) | Lignin | 0.0010 | 1.000 |
| **Ethanone, 1-(3,4-dimethoxyphenyl)-** | **Lignin** | **0.2578** | **0.057** |
| Benzoic acid, 4-hydroxy-3-methoxy-, methyl ester (Vanillic Acid, methyl est" | Lignin | 0.1083 | 0.288 |
| Benzene, 4-ethenyl-1,2-dimethoxy- | Lignin | 0.1627 | 0.256 |
| Benzene, 1-methoxy-4-methyl- | Lignin | 0.1236 | 0.257 |
| 3,4-Dimethoxytoluene | Lignin | 0.1218 | 0.318 |
| Benzene, 1-ethenyl-4-methoxy- | Lignin | 0.1784 | 0.120 |

| | | | |
|---|---|---|---|
| Hexadecanoic acid, methyl ester (Palmitic acid-C16) | Lipid | 0.1797 | 0.129 |
| n-Docosane (C22) | Lipid | 0.1627 | 0.256 |
| 1-Butyne, 3,3-dimethyl- | Lipid | 0.0429 | 0.641 |
| 3-Decene | Lipid | 0.0463 | 0.619 |
| 7-Tetradecene | Lipid | 0.0034 | 0.968 |
| Propene | Lipid | 0.0181 | 0.824 |
| Isocrotonic acid | Lipid | 0.0367 | 0.670 |
| 2-Butenedioic acid, 2-methyl-, (E)- | Lipid | 0.0363 | 0.605 |
| 1-Hexene, 3-methyl- | Lipid | 0.0053 | 0.953 |
| 1-Heptene | Lipid | 0.1226 | 0.258 |
| n-Nonane | Lipid | 0.0633 | 0.496 |
| 1,3-Butadiene | Lipid | 0.0320 | 0.714 |
| C14_alkene_3 | Lipid | 0.0840 | 0.436 |
| 1-Pentene, 3-ethyl-2-methyl- | Lipid | 0.0465 | 0.614 |
| n-Undecane | Lipid | 0.0391 | 0.662 |
| n-Pentadecane | Lipid | 0.0132 | 0.861 |
| 1,3-Octadiene | Lipid | 0.0245 | 0.792 |
| n-Octane | Lipid | 0.1149 | 0.257 |
| n-Heptane | Lipid | 0.1238 | 0.274 |
| n-Tetracosane | Lipid | 0.1627 | 0.256 |
| 1-Hexadecene | Lipid | 0.0507 | 0.591 |
| n-Dodecane | Lipid | 0.0145 | 0.855 |
| Hex-2-yn-4-one, 2-methyl- | Lipid | 0.0376 | 0.678 |
| n-Nonacosane (C29) | Lipid | 0.1134 | 0.285 |
| n-Heptacosane (C27) | Lipid | 0.0913 | 0.361 |
| n-Tridecane | Lipid | 0.0070 | 0.933 |
| n-Heptadecane | Lipid | 0.0160 | 0.832 |
| **n-Decane** | **Lipid** | **0.3689** | **0.006** |
| n-Hexadecane | Lipid | 0.0105 | 0.966 |
| n-Tricosane (C23) | Lipid | 0.0269 | 0.761 |
| n-Tetradecane | Lipid | 0.0470 | 0.621 |
| n-Hexacosane (C26) | Lipid | 0.0232 | 0.816 |
| Hepta-2,4-dienoic acid, methyl ester | Lipid | 0.0454 | 0.639 |
| n-Heneicosane (C21) | Lipid | 0.0339 | 0.703 |
| **Pyrazolo[5,1-c][1,2,4]benzotriazin-8-ol** | **N-Bearing** | **0.3236** | **0.015** |
| N-Butyl-tert-butylamine | N-Bearing | 0.0859 | 0.382 |
| 1H-Tetrazole, 1-methyl- | N-Bearing | 0.0870 | 0.368 |
| **3-Pyridinol** | **N-Bearing** | **0.2657** | **0.025** |
| 1H-Pyrrole-2-carboxaldehyde | N-Bearing | 0.0401 | 0.679 |
| 2,5-Furandione, 3-methyl- | N-Bearing | 0.0473 | 0.638 |
| Aniline | N-Bearing | 0.0364 | 0.683 |
| 1H-Pyrrole, 3-methyl- | N-Bearing | 0.1556 | 0.198 |
| Pyridine 3-methyl | N-Bearing | 0.0757 | 0.414 |
| Pyridine, 4-methoxy- | N-Bearing | 0.0543 | 0.555 |
| p-Aminotoluene | N-Bearing | 0.1208 | 0.264 |
| Hexanedinitrile | N-Bearing | 0.0607 | 0.520 |
| 1H-Pyrrole-2-carboxaldehyde, 1-methyl- | N-Bearing | 0.0578 | 0.543 |
| Pyridine, 2-ethyl | N-Bearing | 0.0407 | 0.650 |
| 3-Methylpyridazine | N-Bearing | 0.0701 | 0.487 |
| 3-Phenylpyridine | N-Bearing | 0.1260 | 0.244 |
| Propane, 2-nitro- | N-Bearing | 0.0112 | 0.892 |
| 4-Pyridinecarboxaldehyde | N-Bearing | 0.0051 | 0.944 |
| 2-Pyridinealdehyde | N-Bearing | 0.0442 | 0.625 |
| 4(1H)-Pyridinone, 2,3-dihydro-1-methyl- | N-Bearing | 0.0157 | 0.853 |
| Alpha-amino-gamma-butyrolactone | N-Bearing | 0.0123 | 0.894 |
| Pyrimidine | N-Bearing | 0.0142 | 0.877 |

| | | | |
|---|---|---|---|
| Piperidine-2,5-dione | N-Bearing | 0.1279 | 0.243 |
| Acetamide, N-(2,4-dihydroxyphenyl)- | N-Bearing | 0.0646 | 0.502 |
| **4-Pyridinecarbonitrile** | **N-Bearing** | **0.3241** | **0.009** |
| Diethyltoluamide (DEET) | N-Bearing | 0.1416 | 0.219 |
| 5H-1-Pyrindine | N-Bearing | 0.0481 | 0.601 |
| 1,4-Benzenediamine | N-Bearing | 0.0104 | 0.883 |
| 5-Dimethylaminopyrimidine | N-Bearing | 0.0358 | 0.699 |
| Ethanone, 1-(1-methyl-1H-pyrrol-2-yl)- | N-Bearing | 0.0464 | 0.626 |
| Acetamide, N-hydroxy | N-Bearing | 0.0332 | 0.762 |
| 2-(N-Methyl-N-ethylamino)phenol | N-Bearing | 0.0626 | 0.507 |
| Cyanamide, dimethyl- | N-Bearing | 0.0339 | 0.703 |
| 2-Pyrimidinamine | N-Bearing | 0.0339 | 0.703 |
| Methylaminoacetonitrile | N-Bearing | 0.0305 | 0.777 |
| 2-Amino-4-methylpyrimidine | N-Bearing | 0.0580 | 0.552 |
| 3-Pyridinecarbonitrile | N-Bearing | 0.0246 | 0.787 |
| 4-Amino-2(1H)-pyridinone | N-Bearing | 0.0332 | 0.762 |
| N-Isopropyl-4-piperidone | N-Bearing | 0.0332 | 0.762 |
| Nicotinyl Alcohol | N-Bearing | 0.0332 | 0.762 |
| Phenol | Phenol | 0.1662 | 0.163 |
| Phenol, 4-methyl- | Phenol | 0.1311 | 0.216 |
| **Benzene, 1,4-dimethoxy-** | **Phenol** | **0.2542** | **0.007** |
| Benzene, 1-ethyl-4-methoxy- | Phenol | 0.0557 | 0.555 |
| 3-Furaldehyde | Polysaccharide | 0.0596 | 0.517 |
| Levoglucosan | Polysaccharide | 0.0194 | 0.830 |
| 2(3H)-Furanone, 5-methyl- | Polysaccharide | 0.0578 | 0.503 |
| 2-Cyclopenten-1-one, 2-hydroxy-3-methyl- | Polysaccharide | 0.1278 | 0.239 |
| Furfural, 5-methyl- | Polysaccharide | 0.0602 | 0.511 |
| Benzofuran, 2,3-dihydro- | Polysaccharide | 0.1360 | 0.239 |
| Acetic anhydride | Polysaccharide | 0.0424 | 0.645 |
| 4H-Pyran-4-one, 3-hydroxy-2-methyl- | Polysaccharide | 0.0013 | 0.982 |
| 2-Furanmethanol | Polysaccharide | 0.0123 | 0.894 |
| Furan, 2,4-dimethyl- | Polysaccharide | 0.0030 | 0.979 |
| 2-Cyclopenten-1-one, 2-methyl- | Polysaccharide | 0.0001 | 1.000 |
| 2-Acetylfuran | Polysaccharide | 0.0519 | 0.558 |
| 2H-Pyran-2-one | Polysaccharide | 0.0598 | 0.510 |
| 1,2-Cyclohexanedione | Polysaccharide | 0.0201 | 0.819 |
| Butanal, 2-methyl- | Polysaccharide | 0.0304 | 0.727 |
| 2-Cyclopenten-1-one, 2,3-dimethyl- | Polysaccharide | 0.0441 | 0.629 |
| Furan, 2,5-dimethyl- | Polysaccharide | 0.0458 | 0.616 |
| Levoglucosenone | Polysaccharide | 0.0826 | 0.398 |
| 2(5H)-Furanone, 5-methyl- | Polysaccharide | 0.0341 | 0.689 |
| Furan, 2-ethyl-5-methyl- | Polysaccharide | 0.0651 | 0.473 |
| Benzofuran, 2-methyl- | Polysaccharide | 0.0596 | 0.501 |
| Furan, 2,3,5-trimethyl- | Polysaccharide | 0.0297 | 0.732 |
| Vinylfuran | Polysaccharide | 0.0928 | 0.342 |
| Cyclopent-2-ene-1-one, 2,3,4-trimethyl- | Polysaccharide | 0.1078 | 0.308 |
| 1,2-Cyclopentanedione, 3-methyl- | Polysaccharide | 0.0332 | 0.762 |
| PENTANAL | Polysaccharide | 0.1613 | 0.162 |
| Furan, 2-ethyl- | Polysaccharide | 0.0826 | 0.432 |
| 1,2-Cyclopentanedione | Polysaccharide | 0.0821 | 0.396 |
| Cyclopropanecarboxaldehyde, methylene- | Polysaccharide | 0.0075 | 0.962 |
| Cyclopentanone | Polysaccharide | 0.0216 | 0.795 |
| 3-Acetamidofuran | Polysaccharide | 0.0331 | 0.698 |
| Furfural | Polysaccharide | 0.0633 | 0.553 |
| 5-Ethyl-2-furaldehyde | Polysaccharide | 0.0348 | 0.719 |
| 2(3H)-Benzofuranone, 3-methyl- | Polysaccharide | 0.0363 | 0.605 |

| | | | |
|---|---|---|---|
| Indole | Protein | 0.1918 | 0.103 |
| Styrene | Protein | 0.0188 | 0.824 |
| Benzyl nitrile | Protein | 0.0403 | 0.659 |
| 3-Methylindole | Protein | 0.1415 | 0.209 |
| Ethylbenzene | Protein | 0.0736 | 0.418 |
| Pyridine | Protein | 0.0587 | 0.543 |
| 4-Pyridinamine | Protein | 0.0503 | 0.586 |
| 1H-Pyrrole, 1-methyl- | Protein | 0.1148 | 0.315 |
| Pyridine, 3,5-dimethyl- | Protein | 0.0250 | 0.758 |
| Pyrrole | Protein | 0.0858 | 0.395 |
| Benzonitrile | Protein | 0.0218 | 0.769 |
| 1 H-Pyrrole, 2-ethyl- | Protein | 0.0911 | 0.358 |
| Pyridine | Protein | 0.0058 | 0.941 |
| Benzenepropanenitrile | Protein | 0.1059 | 0.362 |
| Mequinol | Unspecified | 0.0643 | 0.494 |
| Spiro[2.4]hepta-4,6-diene | Unspecified | 0.0407 | 0.615 |
| **(ISTD) 3-Ethoxy-4-methoxybenzaldehyde** | **Unspecified** | **0.2108** | **0.045** |
| Hydroquinone | Unspecified | 0.0917 | 0.348 |
| 1,3,5-Cycloheptatriene | Unspecified | 0.1108 | 0.284 |
| Methanesulfonic acid, methyl ester | Unspecified | 0.0522 | 0.566 |
| Benzene, 1,4-dimethoxy-2,3,5,6-tetramethyl- | Unspecified | 0.0123 | 0.894 |
| Pyruvaldehyde | Unspecified | 0.1073 | 0.294 |
| Monobenzone | Unspecified | 0.1077 | 0.329 |
| C8_H16 | Unspecified | 0.0498 | 0.594 |
| 1-Undecanol | Unspecified | 0.0118 | 0.882 |
| Indane | Unspecified | 0.0480 | 0.615 |
| Dimethylbenzofuran | Unspecified | 0.0154 | 0.833 |
| 1H-Inden-1-one, 2,3-dihydro- | Unspecified | 0.0081 | 0.905 |
| Phenanthrene | Unspecified | 0.0281 | 0.763 |
| 2,3,6-Trimethylnaphthalene | Unspecified | 0.0782 | 0.436 |
| **Trimethylphenol** | **Unspecified** | **0.3823** | **0.003** |
| Vinyl crotonate | Unspecified | 0.0332 | 0.762 |
| C9_H8 | Unspecified | 0.0029 | 0.969 |
| Toluene | Unspecified | 0.0054 | 0.938 |
| 3-Penten-2-one, (E)- | Unspecified | 0.0091 | 0.906 |
| 4-Hydroxy-2-methylacetophenone | Unspecified | 0.0147 | 0.866 |
| **Benzoic acid, 4-hydroxy-3,5-dimethoxy-, hydrazide** | **Unspecified** | **0.3682** | **0.005** |
| 1-Propene, 3-azido- | Unspecified | 0.0059 | 0.957 |
| **2-Pentanone** | **Unspecified** | **0.2108** | **0.045** |
| Resorcinol (Dihydroxybenzene) | Unspecified | 0.0007 | 0.993 |
| Acenaphthene | Unspecified | 0.0157 | 0.848 |
| Ethanone, 1-(3-hydroxy-4-methoxyphenyl)- | Unspecified | 0.1711 | 0.197 |
| 2-Propenoic acid, ethenyl ester | Unspecified | 0.0197 | 0.803 |
| **2-Methoxy-5-methylphenol** | **Unspecified** | **0.2108** | **0.045** |
| 2-Propen-1-ol | Unspecified | 0.0010 | 1.000 |
| C11_H12 | Unspecified | 0.1354 | 0.201 |
| (E)-1,3-Butadien-1-ol | Unspecified | 0.0553 | 0.574 |
| Thiophene | Unspecified | 0.0123 | 0.894 |
| Benzene, heptyl- | Unspecified | 0.0140 | 0.869 |
| 3-Methylthiophene-2-carbonitrile | Unspecified | 0.1349 | 0.194 |
| Fluoranthene | Unspecified | 0.0296 | 0.745 |
| Pyrene | Unspecified | 0.0181 | 0.823 |
| 7-Methylindan-1-one | Unspecified | 0.0643 | 0.561 |
| Acenaphthylene | Unspecified | 0.0010 | 1.000 |
| **Cyclobut-1-enylmethanol** | **Unspecified** | **0.2108** | **0.045** |
| **2-Methylthiophene** | **Unspecified** | **0.2108** | **0.045** |

| | | | |
|---|---|---|---|
| Isobutyl nitrite | Unspecified | 0.0123 | 0.894 |
| Phenol, 2-methoxy-5-(1-propenyl)-, (E)- | Unspecified | 0.0363 | 0.605 |

**Table S17.** Two-way ANOVA results for the effect of Decomposition Stage, Treatment, and Decomposition Stage x Treatment interaction on molecular richness (top) and molecular evenness (bottom) diversity metrics. Significant P-values ($P \leq 0.1$) are presented in bold.

| Diversity Metric | Factor | Numerator df | Denominator df | F-value | P-value |
|---|---|---|---|---|---|
| Molecular Richness | Decomp Stage | 2 | 69 | 9.4898 | **<0.001** |
| | Treatment | 1 | 70 | 0.0016 | 0.9686 |
| | Decomp Stage x Treatment | 2 | 69 | 0.3979 | 0.6733 |
| Molecular Evenness | Decomp Stage | 2 | 69 | 12.224 | **<0.001** |
| | Treatment | 1 | 70 | 0.3925 | 0.5331 |
| | Decomp Stage x Treatment | 2 | 69 | 0.6333 | 0.5340 |

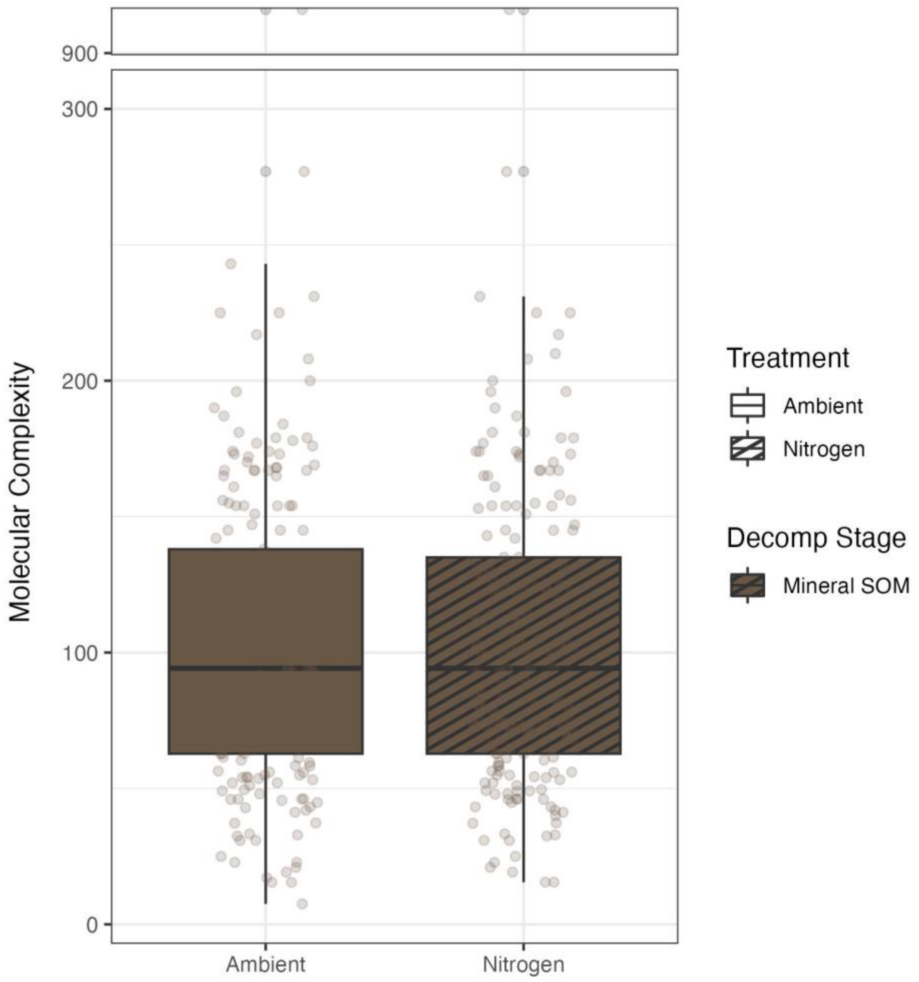


**Figure S5.** The effect of the nitrogen deposition treatment on the molecular complexity of individual biochemical compounds present in the mineral SOM (*n* = 386). No treatment effect was detected by ANOVA ($F_{1,384} = 0$, $P = 0.997$).

**Table S18.** Tukey's HSD pairwise comparisons where two-way ANOVA of molecular diversity metrics revealed significant ($P \leq 0.1$) effect of Decomposition Stage on molecular richness (top) and evenness (bottom) (see Supplementary Table S17). Significant P-values ($P \leq 0.1$) are presented in bold.

| Diversity Metric | Factor | Post-hoc Comparison | Mean Difference | P-value |
|---|---|---|---|---|
| Molecular Richness | Decomp Stage | Undecomposed – Decomposed | -12.8750 | **0.0011** |
| | | Decomposed – Mineral SOM | 13.0833 | **<0.001** |
| | | Undecomposed – Mineral SOM | 0.2083 | 0.9980 |
| | Decomp Stage x Treatment | Ambient Undecomposed – Ambient Decomposed | -12.5833 | **0.0316** |
| | | Ambient Decomposed – Ambient Mineral SOM | 3.0000 | 0.8116 |
| | | Ambient Undecomposed – Ambient Mineral SOM | 15.5833 | **0.0059** |
| | | Nitrogen Undecomposed – Nitrogen Decomposed | -13.1667 | **0.0232** |
| | | Nitrogen Decomposed – Nitrogen Mineral SOM | 10.5834 | **0.0830** |
| | | Nitrogen Undecomposed – Nitrogen Mineral SOM | -2.5833 | 0.8565 |
| Molecular Evenness | Decomp Stage | Undecomposed – Decomposed | -0.0580 | **0.0026** |
| | | Decomposed – Mineral SOM | -0.0220 | 0.3902 |
| | | Undecomposed – Mineral SOM | -0.0800 | **<0.001** |
| | Decomp Stage x Treatment | Ambient Undecomposed – Ambient Decomposed | -0.0624 | **0.0274** |
| | | Ambient Decomposed – Ambient Mineral SOM | -0.0040 | 0.9843 |
| | | Ambient Undecomposed – Ambient Mineral SOM | -0.0664 | **0.0176** |
| | | Nitrogen Undecomposed – Nitrogen Decomposed | -0.0536 | **0.0673** |
| | | Nitrogen Decomposed – Nitrogen Mineral SOM | -0.0401 | 0.2136 |
| | | Nitrogen Undecomposed – Nitrogen Mineral SOM | -0.0937 | **0.0005** |

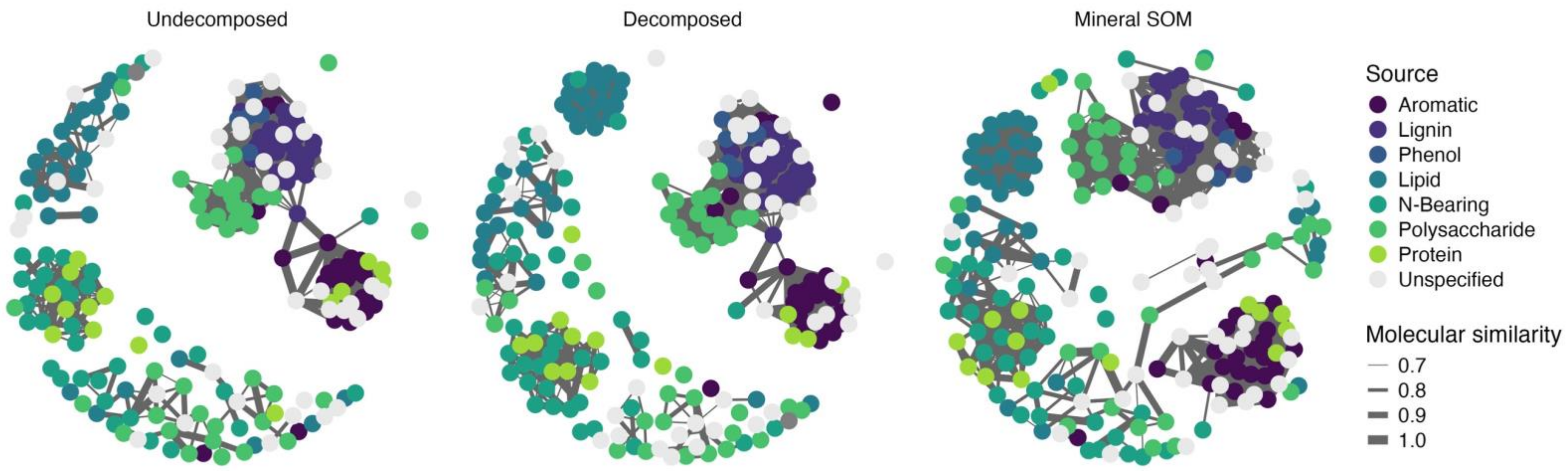


**Figure S6.** Molecular network showing relationships between the biochemical compounds at each decomposition stage (*n* = 3). Nodes represent individual molecules (Undecomposed: *n* = 209; Decomposed: *n* = 225, Mineral SOM: *n* = 216) and edges represent the quantity of interactions and relationships between those molecules (Undecomposed: *n* = 2950; Decomposed: *n* = 3372, Mineral SOM: *n* = 3020). Thickness of the edges is determined by the molecular similarity of which the cut off value was set to 0.7. Nodes are colored by their relative abundance.

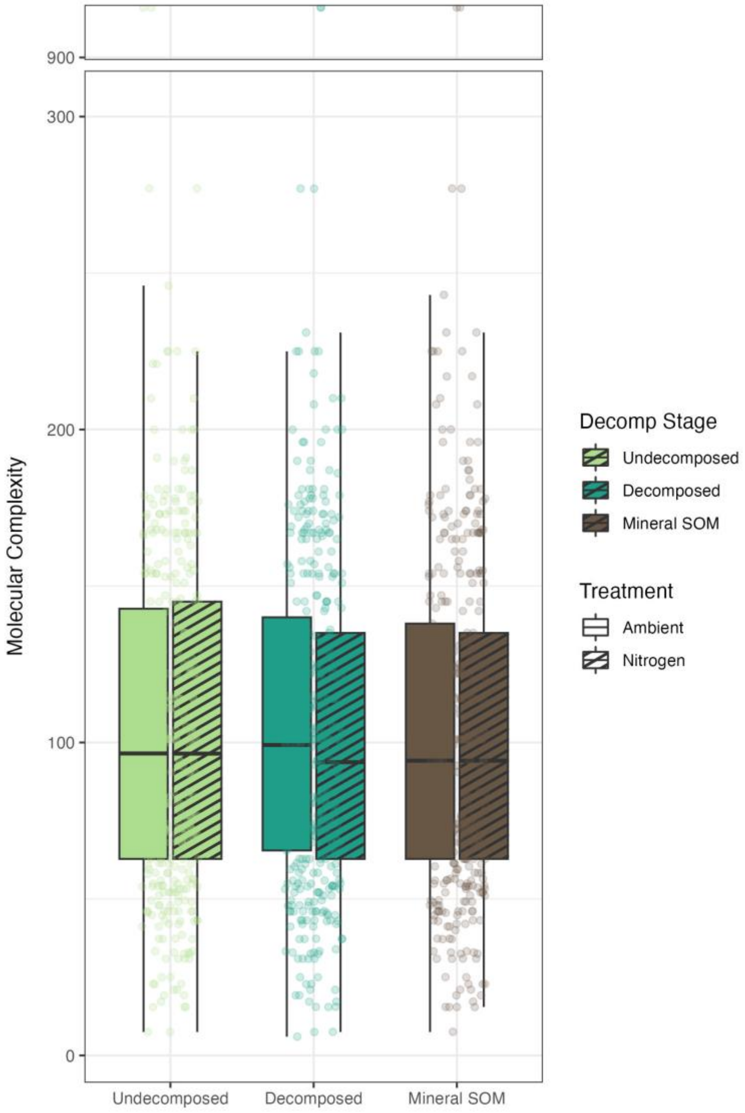


**Figure S7.** The effect of decomposition and the nitrogen deposition treatment on the molecular complexity of individual biochemical compounds present at each decomposition stage ($n_{\text{total}}$ = 1182). ANOVA detected no difference in decomposition stage (F = 0.01, $P$ = 0.989), treatment (F = 0.03, $P$ = 0.852), or their interaction (F = 0.02, $P$ = 0.976).
